# Multivalent interactions drive the *Toxoplasma* AC9:AC10:ERK7 complex to concentrate ERK7 in the apical cap

**DOI:** 10.1101/2022.01.04.474968

**Authors:** Peter S. Back, William J. O’Shaughnessy, Andy S. Moon, Pravin S. Dewangan, Michael L. Reese, Peter J. Bradley

## Abstract

The *Toxoplasma* inner membrane complex (IMC) is a specialized organelle that is crucial for the parasite to establish an intracellular lifestyle and ultimately cause disease. The IMC is composed of both membrane and cytoskeletal components, further delineated into the apical cap, body, and basal subcompartments. The apical cap cytoskeleton was recently demonstrated to govern the stability of the apical complex, which controls parasite motility, invasion, and egress. While this role was determined by individually assessing the apical cap proteins AC9, AC10, and the MAP kinase ERK7, how the three proteins collaborate to stabilize the apical complex is unknown. In this study, we use a combination of deletion analyses and yeast-2-hybrid experiments to establish that these proteins form an essential complex in the apical cap. We show that AC10 is a foundational component of the AC10:AC9:ERK7 complex and demonstrate that the interactions among them are critical to maintain the apical complex. Importantly, we identify multiple independent regions of pairwise interaction between each of the three proteins, suggesting that the AC9:AC10:ERK7 complex is organized by multivalent interactions. Together, these data support a model in which multiple interacting domains enable the oligomerization of the AC9:AC10:ERK7 complex and its assembly into the cytoskeletal IMC, which serves as a structural scaffold that concentrates ERK7 kinase activity in the apical cap.

**IMPORTANCE:** The phylum Apicomplexa consists of obligate, intracellular parasites including the causative agents of toxoplasmosis, malaria, and cryptosporidiosis. Hallmarks of these parasites are the IMC and the apical complex, both of which are unique structures that are conserved throughout the phylum and required for parasite survival. The apical cap portion of the IMC has previously been shown to stabilize the apical complex. Here, we expand on those studies to determine the precise protein-protein interactions of the apical cap complex that confer this essential function. We describe the multivalent nature of these interactions and show that the resulting protein oligomers likely tether ERK7 in the apical cap. This study represents the first description of the architecture of the apical cap at a molecular level, expanding our understanding of the unique cell biology that drives *Toxoplasma* infections.

## INTRODUCTION

The phylum Apicomplexa contains a large group of obligate intracellular parasites of medical and veterinary importance (1). Human parasites include *Toxoplasma gondii*, which causes toxoplasmosis in immunocompromised people and congenitally infected neonates; *Plasmodium* spp., which causes malaria; and *Cryptosporidium* spp., which causes diarrheal disease in children (2–4). Important animal pathogens include *Neospora* spp., *Eimeria* spp., *Theileria* spp., and *Babesia* spp., which together account for enormous economic losses in the poultry and cattle industries (5–7). These apicomplexan parasites require specialized machinery to actively invade their mammalian host cells, establish an intracellular niche, and cause disease. The alveoli are one such structure and are formed from a series of flattened membranous vesicles that underlies the plasma membrane. The alveoli represent a hallmark of the broader superphylum Alveolata that includes ciliates, dinoflagellates, and apicomplexan parasites (8).

In apicomplexans, the alveoli are called the inner membrane complex (IMC). The IMC is a peripheral membrane system with two well described roles: a platform to anchor the glideosome, the actin-myosin motor complex that interacts with micronemal adhesins secreted onto the parasite surface for gliding motility, and a scaffold for endodyogeny, an internal budding process of replication (9, 10). The IMC is situated between the plasma membrane and cortical microtubules at the periphery of the cell and consists of a series of flattened membrane vesicles and an underlying cytoskeletal network of intermediate filament-like proteins called the alveolins (11, 12). The membrane vesicles are organized into rectangular plates along the body of the parasite, culminating in a single cone-shaped plate at the apex called the apical cap (13, 14). Because both the apical cap and body sections of the IMC are composed of similar membrane and cytoskeletal components, they were previously believed to be one unified structure. However, the discovery of an array of new IMC proteins revealed that the apical cap contains a unique cohort of proteins, suggesting a specialized function for this region (15–20). Recent analyses of a group of these proteins revealed a third IMC function – regulating the biogenesis and stability of the apical complex (21–23).

The apical complex is a group of cytoskeletal structures at the apex of the parasite that includes the microtubule-based conoid, the flanking apical polar ring (APR), and two preconoidal rings (19, 24, 25). The striking basket-shaped ultrastructure of the conoid allowed it to be readily described in the tissue cyst-forming coccidian subgroup of the Apicomplexa (e.g. *Toxoplasma*, *Sarcocystis, Eimeria*). Remarkably, the apical complex, including the conoid, has been described in early-branching alveolates that are not members of Apicomplexa, suggesting the structure is more ancient than originally appreciated (26, 27). Indeed, while the conoid was originally presumed to be missing from Haemosporidia (1, 28), recent studies have identified a reduced conoid complex in multiple stages of *Plasmodium*, suggesting that this structure is conserved throughout the Apicomplexa (29–31). Moreover, the apical complex contains orthologs of cilium-associated proteins, leading to a potential link between the apical complex of apicomplexan parasites and more typical eukaryotic cilia (29, 32–35). Numerous studies have demonstrated that the apical complex regulates the secretion of specialized organelles called micronemes and rhoptries, which govern parasite motility, attachment, invasion, and egress (36). While the trigger for rhoptry secretion at the apical complex is unknown, calcium signaling cascades have been shown to coordinate both microneme secretion and conoid extrusion, suggesting a connection between the two activities (37). The conoid has also been implicated in initiating motility via several calmodulin-like proteins, the myosin motor protein MyoH, and the essential formin protein FRM1 (38–40). In addition, several APR-localizing proteins were shown to be important in controlling microneme release, indicating that these flanking cytoskeletal structures also contribute to the function of the apical complex (19, 41, 42).

While the molecular composition and function of the apical complex is becoming clearer, how it is formed and maintained is largely a mystery. Recently, three apical cap proteins (AC9, AC10, and ERK7) were identified as essential for the maturation of the apical complex (21–23). Depleting any one of these proteins eliminates the conoid in mature parasites, resulting in a complete block in motility, invasion, and egress. Importantly, AC9 was shown to accomplish this by recruiting the conserved MAP kinase ERK7 to the apical cap and regulating its kinase activity (23). Thus, it is evident that AC9, AC10, and ERK7 work in conjunction to facilitate the apical complex maturation and function. However, how these proteins interact and coordinate at the apical cap to confer their functions remains unknown. In this study, we explore the organization and mechanism of this essential protein complex. We show that AC10 recruits both AC9 and ERK7 to the apical cap, suggesting it is the anchor for the complex. We combine yeast-2-hybrid experiments to examine direct pairwise interactions with deletion analyses in parasites to assess the functional importance of these interactions. Through these experiments, we reveal multiple domains in AC9 and AC10 that are critical for assembling the complex at the apical cap and for the maturation of the conoid. Importantly, we show that these domains mediate independent pairwise interactions between AC9, AC10, and ERK7. Thus, we propose that these multimeric interactions drive the oligomerization of the AC9:AC10:ERK7 complex into the apical cap cytoskeleton, which tethers ERK7 to the site of its essential function in coordinating the proper biogenesis of the apical complex.

## RESULTS

### AC10 is essential for recruitment of the AC9:AC10:ERK7 complex to the apical cap

While AC9, AC10 and ERK7 were recently shown to be essential for apical complex assembly and stabilization (21–23), the interactions between the three proteins and how they are organized in the apical cap remain poorly understood (an overview of these proteins is shown in Fig. 1). To explore their interactions, we generated parasites with AC10 tagged with an auxin-inducible degron fused to 3xHA, AC9 tagged with 3xMyc, and ERK7 tagged with 3xTy (triple-tagged: AC10^AID-3xHA^/AC9^3xMyc^/ERK7^3xTy^). As shown previously, the AC10^AID-3xHA^ fusion protein targets correctly to the apical cap, degrades efficiently upon addition of auxin (IAA), and results in the loss of AC9 from the apical cap (Fig. 2A and B) (22). Our triple-tagged parasites allowed us to additionally demonstrate that AC10^AID-3xHA^ knockdown removes ERK7 from the apical cap though its cytoplasmic staining is retained (Fig. 2B). We used line intensity scans to quantify the levels of ERK7 at the apical cap versus the bulk cytosol, which clearly demonstrated a loss in concentrated apical cap signal upon AC10 knockdown (Fig. S2). Consistent with the AC9 and ERK7 staining patterns, western blot analyses showed that AC9 is predominantly degraded while ERK7 levels appear to remain stable (Fig. 2C) (22). In agreement with previous studies (22), depletion of AC10 results in the elimination of the conoid (Fig. 2D), which is lethal for the parasites (Fig. 2E), as it renders them immotile and noninvasive. In addition, we confirmed that the knockdown of AC9 does not affect the localization of AC10 (Fig. 2F) (22), indicating that AC10 does not rely on AC9 for apical cap localization. These results demonstrate that AC10 is essential for recruiting both AC9 and ERK7 to the apical cap and suggest that AC10 is the foundational component of the AC9:AC10:ERK7 complex.

**Fig 1.**
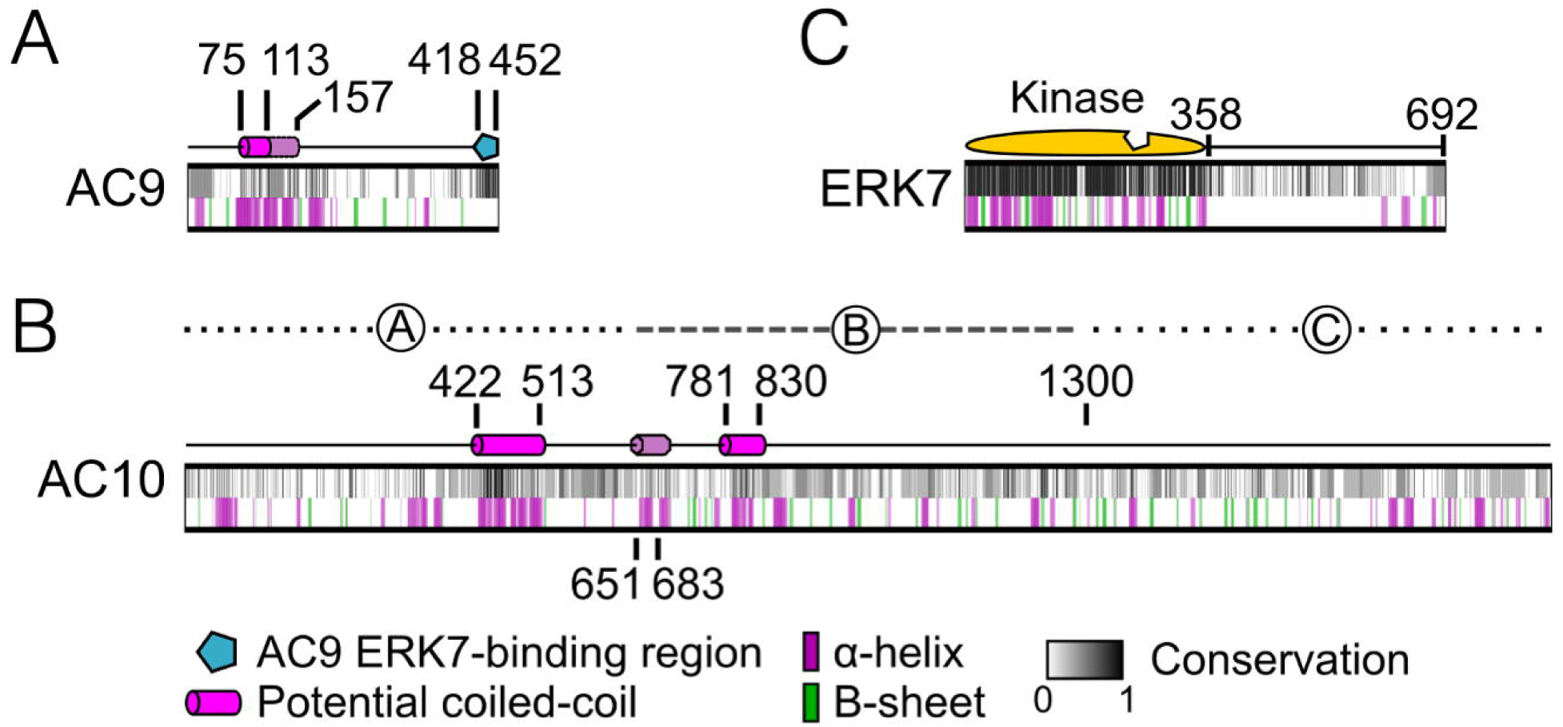
Overview of AC9, AC10, and ERK7 domains. (A) Diagram of AC9 illustrates a predicted coiled-coil (CC) domain (residues 75-113), conserved α-helices flanking the CC domain (residues 113-157), and the ERK7-binding region (residues 418-452). (B) Diagram of AC10 contains two predicted CC domains (CC1: 422-513 and CC2: 781-830) as well as a short conserved α-helix (651-683). Regions A (2-650), B (651-1300), and C (1301-1979) delineate the divisions of AC10 used for yeast-2-hybrid (Y2H) assays. (C) Diagram of ERK7 showing the kinase domain (1-358) including the active site (notched region) and the C-terminus (359-692). All three diagrams contain a grayscale representation of the degree of conservation as well as secondary structure predictions which are depicted by purple and green bars as noted in the legend. Conservation calculations are based on multiple sequence alignments of AC9, AC10, and ERK7 sequences from *T. gondii, N. caninum, B. besnoitia, C. suis, E. maxima,* and *E. tenella*.

**Fig 2.**
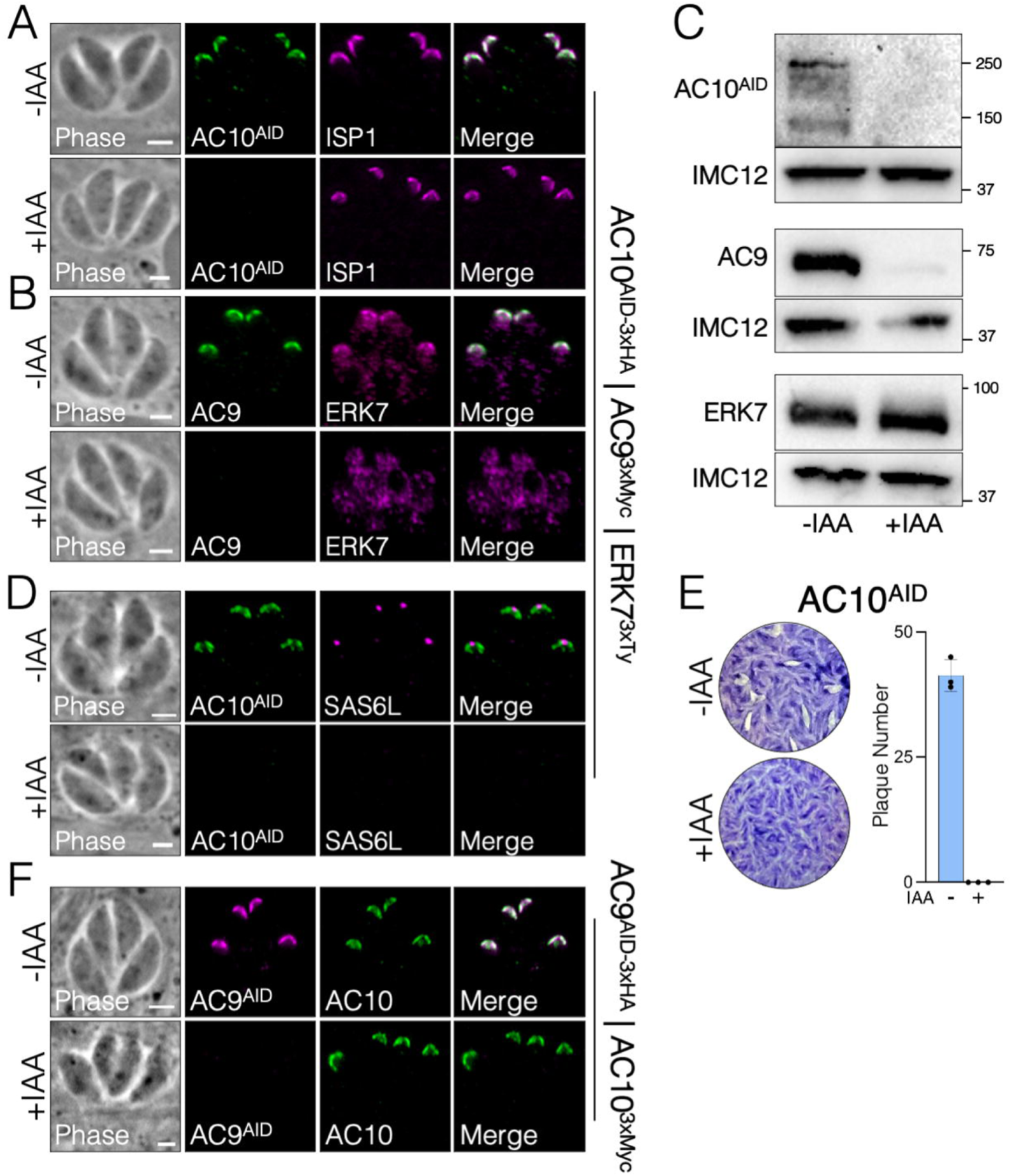
AC10 is an essential component of the apical cap. (A) Immunofluorescence assay (IFA) of triple-tagged parasites (AC10^AID-3xHA^ | AC9^3xMyc^ | ERK7^3xTy^) shows that endogenous AC10^AID-3xHA^ co-localizes with the apical cap marker ISP1 and is efficiently depleted upon addition of IAA (3-indoleacetic acid). Green, rabbit anti-HA; magenta, mouse anti-ISP1. (B) IFA showing that the depletion of AC10^AID-3xHA^ results in the absence of AC9 and the loss of ERK7 from the apical cap. Green, rabbit anti-Myc; magenta, mouse anti-Ty. (C) Western blot analysis confirms efficient degradation of AC10^AID-3xHA^ and the concomitant nearly complete degradation of AC9 upon AC10^AID-3xHA^ knockdown. ERK7 levels are not substantially affected. AC10^AID-3xHA^, mouse anti-HA; AC9, mouse anti-Myc; ERK7, mouse anti-Ty. Rabbit anti-IMC12 was used as a loading control and validation of this antibody is shown in Fig. S1. (D) AC10^AID-3xHA^ knockdown results in the elimination of the conoid, detected by SAS6L. Green, rabbit anti-HA; magenta, mouse anti-SAS6L. (E) Representative plaque assay images and quantification of plaque numbers illustrate a complete loss of plaque formation upon AC10^AID-3xHA^ depletion. (F) Using parasites tagged with AC9^AID-3xHA^ and AC10^3xMyc^, IFA shows that conditional knockdown of AC9 (+IAA) does not affect the localization of AC10. Green, mouse anti-Myc; magenta, rabbit anti-HA. All scale bars are 2 µm.

### AC9 is recruited to the apical cap through a direct interaction with AC10

Like most IMC components, AC9 and AC10 lack significant homology to other proteins. Both proteins contain large stretches of predicted intrinsic disorder, as well as predicted coiled-coil (CC) domains towards their N-termini (Fig. 1A and B). In addition, we previously identified a well-conserved sequence in the AC9 C-terminus that is required to recruit ERK7 to the apical cap and acts as a competitive inhibitor of ERK7 kinase activity by occupying both the kinase scaffolding and active sites (23). Since AC10 likely recruits AC9 to the apical cap, we reasoned that the AC9 CC domain may be required for this interaction. In the background of our AC9^AID-3xHA^ strain (23), we expressed a second-copy of AC9 driven by the ISC6 promoter and targeted to the UPRT locus (AC9^wt^, Fig. 3A and B) (43). As expected, expression of AC9^wt^ rescued the AC9^AID-3xHA^ knockdown phenotype, as assessed by SAS6L staining of the conoid and plaque assay (Fig. 3C-E). We also created a strain expressing AC9 in which the core of the predicted CC domain had been deleted (residues Δ75-113, AC9^ΔCC^, Fig. 3F). Consistent with the high conservation of this region (Fig. 1A), AC9^ΔCC^ was not correctly targeted to the apical cap and thus it was unable to rescue the effects of AC9^AID-3xHA^ degradation (Fig. 3G-I). Because AC9^ΔCC^ staining was faint, we assessed its stability by western blot and found that it is expressed at the appropriate size, but its protein level is greatly diminished (Fig. S3A). This low level of AC9^ΔCC^ is likely the result of turnover upon loss of binding to its partner AC10 as loss of AC9 is also seen following AC10^AID^ knockdown (Fig. 2C). While we and others have demonstrated a potential interaction between AC9 and AC10 through proximity biotinylation (22, 23), this interaction may either be direct or through an intermediate protein. To test whether AC9 directly binds AC10, we used a yeast-2-hybrid (Y2H) system in which stable interactions drive the expression of the HIS3 marker. Full-length AC9 was expressed as an N-terminal fusion with the LexA DNA binding domain and AC10 was expressed as an N-terminal fusion with the GAL4 activating domain. As AC10 is a large protein of 1979 residues, we split the protein in thirds and tested each portion for activation: AC10^A^ containing residues 2-650, AC10^B^ containing residues 651-1300, AC10^C^ containing residues 1301-1979 (Fig. 1B). Intriguingly, we found that AC9 interacts with two independent regions of AC10, robustly binding both AC10^A^ and AC10^B^; however, we observed no growth in restrictive conditions with the C-terminal AC10^C^ region (Fig. 3J, an overview of all Y2H data is shown in Table 1). These data suggest AC10^C^ does not bind AC9, though we cannot rule out that AC10^C^ is not stable in yeast and is therefore unavailable for binding.

**Fig 3.**
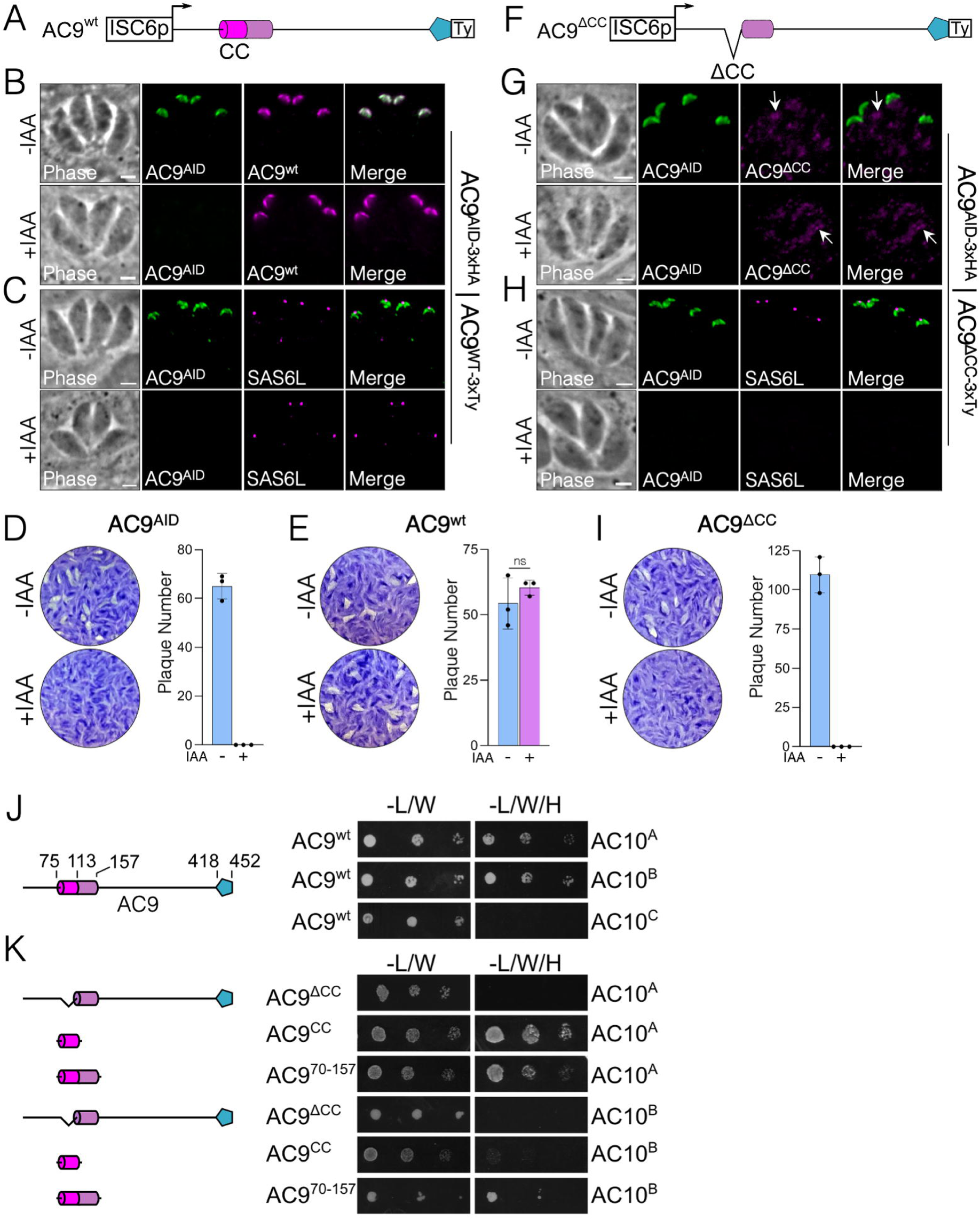
AC9 coiled-coil domain is necessary for localization and function. (A) Diagram of full-length AC9 driven by the ISC6 promoter and a C-terminal 3xTy epitope tag. The AC9 CC domains, α-helices, and the ERK7-binding region are highlighted as in Fig. 1A. (B) IFAs show that the full-length complementation (AC9^wt^) targets correctly to the apical cap and is not affected by the knockdown of endogenous AC9^AID-3xHA^. Green, rabbit anti-HA; magenta, mouse anti-Ty. (C) Staining with SAS6L indicates that the conoid is restored via complementation. Green, rabbit anti-HA; magenta, mouse anti-SAS6L. (D, E) Representative plaque assays and quantification of plaque numbers demonstrate that AC9^AID-3xHA^ depletion results in no plaques while complementation with AC9^wt^ fully restores the plaque defect. (F) Diagram of AC9^ΔCC^ with residues 75-113 deleted from the AC9^wt^ construct. (G) AC9^ΔCC^ fails to localize to the apical cap with faint, dispersed cytoplasmic staining (arrows) upon knockdown of endogenous AC9^AID-3xHA^. Green, rabbit anti-HA; magenta, mouse anti-Ty. (H) As expected from its mislocalization, AC9^ΔCC^ fails to rescue SAS6L staining upon AC9^AID-3xHA^ knockdown. Green, rabbit anti-HA; magenta, mouse anti-SAS6L. All scale bars are 2 µm. (I) Representative plaque assays and their quantifications demonstrate that complementation with AC9^ΔCC^ cannot rescue the plaque defect. (J) Yeast expressing AC9^wt^ and the indicated AC10 constructs were grown in permissive (-L/W) or restrictive (-L/W/H) conditions to assess interaction. A corresponding diagram of full-length AC9 is shown. (K) Y2H assessing the interaction of AC9 mutants with the indicated AC10 sequence, as in (J). Corresponding diagrams of AC9 deletion constructs are shown.

**Table 1:** Overview of yeast-two-hybrid data. Bait and prey constructs and their relative growth on selective media are noted.

| Bait | Prey | Growth |
| --- | --- | --- |
| AC9<sup>wt</sup> | AC10<sup>A</sup> | +++ |
| AC9<sup>wt</sup> | AC10<sup>B</sup> | +++ |
| AC9<sup>wt</sup> | AC10<sup>C</sup> | - |
| AC9<sup>wt</sup> | AC10<sup>AΔCC1</sup> | - |
| AC9<sup>wt</sup> | AC10<sup>AC9BD (651-683)</sup> | +++ |
| AC9<sup>wt</sup> | AC10<sup>684-1300</sup> | ++ |
| AC9<sup>wt</sup> | AC10<sup>684-1300(ΔCC2)</sup> | ++ |
| AC9<sup>ΔCC</sup> | AC10<sup>A</sup> | - |
| AC9<sup>CC</sup> | AC10<sup>A</sup> | + |
| AC9<sup>70-157</sup> | AC10<sup>A</sup> | +++ |
| AC9<sup>ΔCC</sup> | AC10<sup>B</sup> | - |
| AC9<sup>CC</sup> | AC10<sup>B</sup> | + |
| AC9<sup>70-157</sup> | AC10<sup>B</sup> | +++ |
| AC9<sup>wt</sup> | AC10<sup>CC1</sup> | - |
| AC9<sup>wt</sup> | AC10<sup>684-913</sup> | - |
| AC9<sup>wt</sup> | AC10<sup>914-1300</sup> | - |
| ERK7<sup>Kinase</sup> | AC10<sup>A</sup> | +++ |
| ERK7<sup>Kinase</sup> | AC10<sup>B</sup> | - |
| ERK7<sup>Kinase</sup> | AC10<sup>C</sup> | - |
| ERK7<sup>Kinase</sup> | AC10<sup>AΔCC1</sup> | ++ |
| ERK7<sup>C-term</sup> | AC10<sup>A</sup> | - |
| ERK7<sup>C-term</sup> | AC10<sup>B</sup> | ++ |
| ERK7<sup>C-term</sup> | AC10<sup>C</sup> | - |

To test whether the AC9 CC domain was required for this interaction, we deleted this region from the full length Y2H construct (AC9^ΔCC^). Consistent with its inability to rescue the AC9^AID-3xHA^ knockdown phenotype in parasites, AC9^ΔCC^ was unable to bind either AC10^A^ or AC10^B^ (Fig. 3K). Moreover, the AC9 CC domain alone was sufficient to bind AC10^A^ in the Y2H assay, though it could not interact with AC10^B^. The α-helical region of AC9 C-terminal to the predicted CC is one of the more highly conserved areas in the protein (Fig. 1A). We therefore extended our Y2H construct to include this region (AC9^70-157^), which now robustly interacted with both AC10^A^ and AC10^B^ (Fig. 3K). Taken together, these data demonstrate that the conserved α-helical sequence containing the predicted AC9 CC domain is driving interaction with at least two independent sites on AC10, and these interactions are required for forming the functional ternary complex in the apical cap.

### The N-terminal third of AC10 binds both AC9 and ERK7 and is required for efficient recruitment of ERK7 to the apical cap

As AC9^CC^ binds AC10 at multiple distinct sites within the first two thirds of the protein (Fig. 3J), we sought to further delineate which regions of AC10 are required for this interaction. Since AC10^A^ encompasses the most conserved stretch of residues in AC10 and includes a predicted CC domain (Fig. 1B), we generated a Y2H construct in which CC1 was deleted from this region (residues Δ422-513, AC10^A(ΔCC1)^). The Y2H assay showed that AC10^A(ΔCC1)^ was unable to interact with full-length AC9, demonstrating that CC1 is necessary for binding (Fig. 4A). AC10^CC1^ alone was not, however, sufficient to bind AC9, suggesting that this region does not form a simple coiled-coil interaction with AC9 (Fig. 4A).

**Fig 4.**
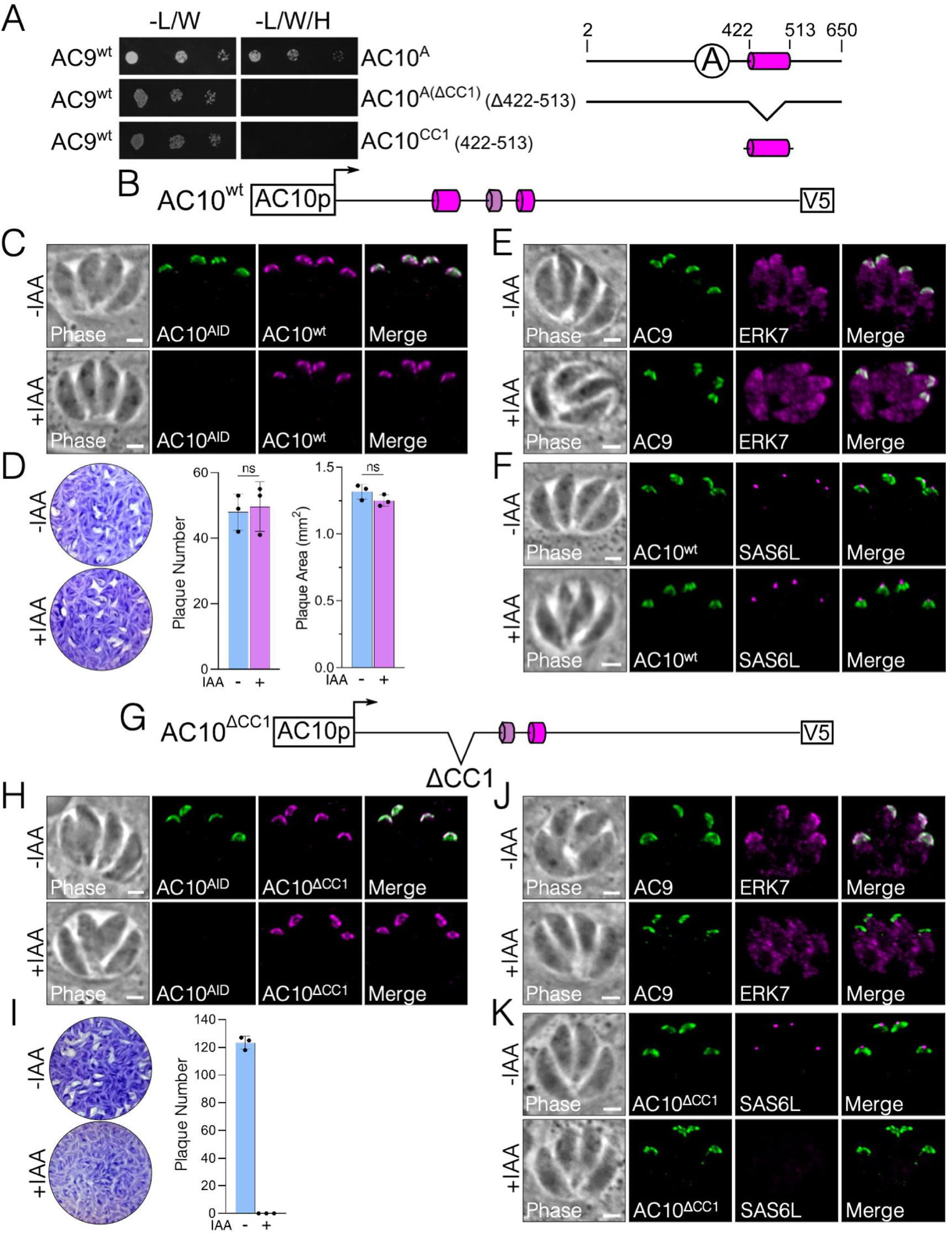
AC10 CC1 binds both AC9 and ERK7 and is essential for apical cap function. (A) Y2H assessing interaction of full-length AC9 with the indicated AC10 constructs, which are shown with corresponding diagrams. The data for AC9:AC10^A^ are shown again from Fig. 3J to facilitate a direct comparison. (B) Diagram of the full-length AC10 with a C-terminal V5 epitope tag (denoted as AC10^wt^). (C) IFA shows that AC10^wt^ localizes properly to the apical cap, which is not affected by knockdown of the endogenous AC10^AID-3xHA^. Green, rabbit anti-HA; magenta, mouse anti-V5. (D) Representative plaque assay images and the corresponding quantification of plaque number and plaque size illustrate that AC10^wt^ fully rescues the lytic ability of AC10^AID-3xHA^ knockdown. Statistical significance was calculated using two-sample two-tailed t tests. (E) IFA demonstrates that AC10^wt^ rescues AC9 and ERK7 localization in the apical cap. Green, rabbit anti-Myc; magenta, mouse anti-Ty. (F) IFA using SAS6L shows that AC10^wt^ restores the conoid +IAA. Green, rabbit anti-V5; magenta, mouse anti-SAS6L. (G) Diagram of AC10^ΔCC1^ with residues 422-513 deleted from the AC10^wt^ construct. (H) IFA shows that AC10^ΔCC1^ targets properly to the apical cap regardless of AC10^AID-3xHA^ knockdown. Green, rabbit anti-HA; magenta, mouse anti-V5. (I) Plaque assays demonstrate that AC10^ΔCC1^ cannot rescue the parasite’s lytic ability. (J) IFA shows that AC9 is present in the apical cap while ERK7 is mislocalized to the cytoplasm upon knockdown of AC10^AID-3xHA^. Green, rabbit anti-Myc; magenta, mouse anti-Ty. (K) IFA illustrates that AC10^ΔCC1^ does not rescue SAS6L localization, indicating the absence of the conoid. Green, rabbit anti-V5; magenta, mouse anti-SAS6L. All scale bars are 2 µm.

To interrogate the functional domains of AC10 in parasites, we expressed full-length AC10 fused to a V5 epitope tag driven by its endogenous promoter and targeted to the UPRT locus (AC10^wt^, Fig. 4B). As expected, the AC10^wt^ complementation construct correctly localized to the apical cap (Fig. 4C), fully rescued the plaque defect (Fig. 4D), properly recruited both AC9 and ERK7 (Fig. 4E), and restored SAS6L staining to the conoid upon AC10^AID-3xHA^ degradation (Fig. 4F). Thus, this complementation system serves as a platform to assess the functional domains of AC10.

To assess the role of AC10^CC1^ in parasites, we deleted CC1 from the full-length construct (AC10^ΔCC1^) and expressed it in the AC10^AID-3xHA^ strain (Fig. 4G). While AC10^ΔCC1^ targeted correctly (Fig. 4H), this complemented strain was unable to form plaques upon AC10^AID-3xHA^ degradation, demonstrating that CC1 is essential for AC10 function (Fig. 4I). Consistent with the lack of plaque formation, AC10^ΔCC1^ did not recruit ERK7 to the apical cap upon AC10^AID-3xHA^ degradation (Fig. 4J), resulting in the loss of SAS6L signal (Fig. 4K). However, we still observed AC9 recruitment in AC10^ΔCC1^ parasites upon AC10^AID-3xHA^ degradation (Fig. 4J). This observation was surprising as we have previously shown that the AC9 C-terminus forms a tight interaction with ERK7 and is required for its recruitment to the apical cap (23). These data suggest that AC10^CC1^ may also directly bind ERK7 independently of the AC10 recruitment of AC9 to the apical cap.

We tested this hypothesis using our Y2H assay and found that AC10^A^ was indeed able to bind the ERK7 kinase domain (Fig. 5). In contrast to the interaction with AC9, in which AC10^CC1^ was required, we found that AC10^A(ΔCC1)^ was still able to bind ERK7 in the Y2H assay, though the interaction was attenuated. In addition to AC10^A^ interacting with the ERK7 kinase domain, we were surprised to find that AC10^B^ also interacted with the intrinsically disordered C-terminus of ERK7, suggesting that ERK7 forms multivalent interactions with AC10. Thus, the Y2H and functional data indicate that multiple AC10 regions mediate interactions with both AC9 (Fig. 3, 4) and ERK7 (Fig. 5). Among these interactions, AC10^CC1^ is required for the efficient recruitment of ERK7 to the apical cap independently of AC9, and this interaction is essential for the formation of the mature conoid.

**Fig 5.**
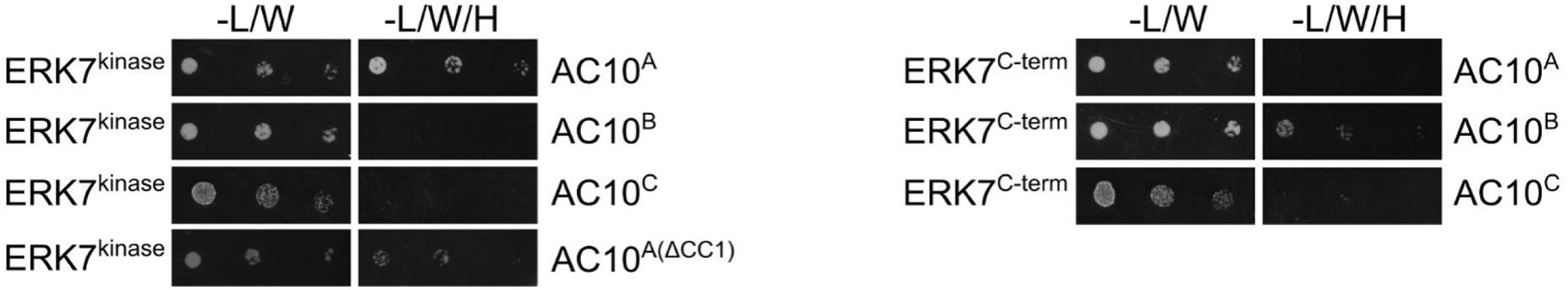
Both regions of ERK7 interact with multiple regions of AC10. Y2H assay to assess interaction of ERK7^kinase^ (2-358) or ERK7^C-term^ (359-652) with the indicated AC10 constructs.

### A short, conserved sequence in AC10 is essential to bind and recruit AC9 to the apical cap

Because the AC10^ΔCC1^ strain was still able to recruit AC9 to the apical cap, we sought to identify additional regions in AC10 that are required for AC9 recruitment. Our Y2H experiments identified regions in AC10^B^ that independently bound AC9 (Fig. 3J). To identify a minimal region that was sufficient for AC9 binding, we focused on a short, conserved sequence within AC10^B^ that is predicted to form an α-helix (Fig. 1B) and has a heptad repeat similar to that seen in coiled-coil domains (Fig. 6A). Y2H analysis showed that residues 651-683 were sufficient to robustly interact with AC9 (Fig. 6B), leading us to label this region as the AC9 binding domain (AC10^AC9-BD^). To test the importance of this region for AC10 function in parasites, we complemented the AC10^AID-3xHA^ strain with a construct in which the AC9-BD had been deleted (AC10^Δ(AC9-BD)^, Fig. 6C). We found that while the truncated protein localized properly to the apical cap (Fig. 6D), it was unable to rescue the plaque defect upon AC10^AID-3xHA^ knockdown (Fig. 6E). We also observed that both AC9 and ERK7 were absent in the apical cap upon AC10^AID-3xHA^ degradation (Fig. 6F), resulting in the loss of the conoid (Fig. 6G). These results suggest that AC10^AC9-BD^ likely forms a short coiled-coil with AC9^CC^, and this interaction is absolutely required for recruitment of the AC9:ERK7 complex to the apical cap in parasites.

**Fig 6.**
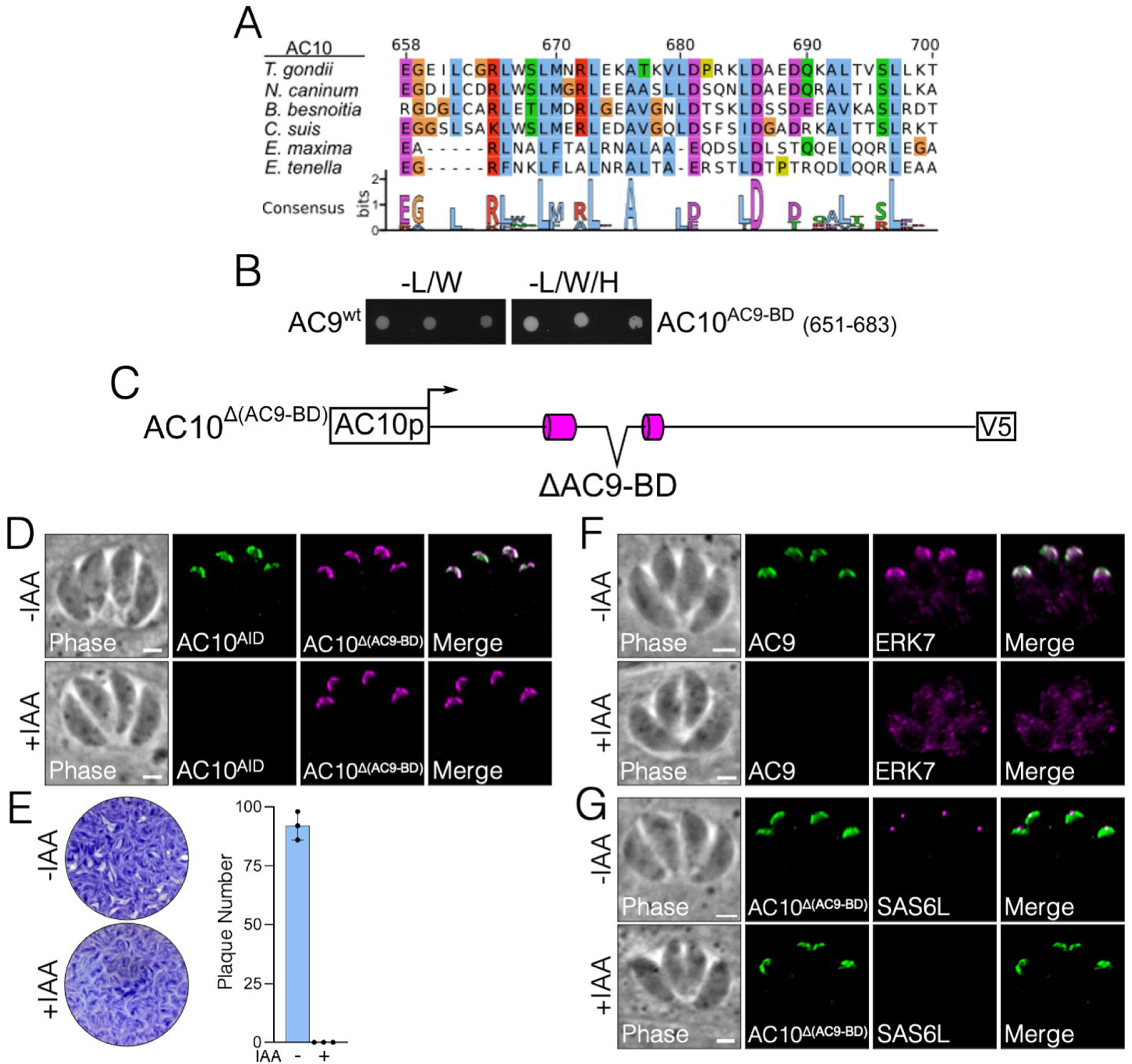
The conserved AC9 binding domain within AC10 is essential for AC10 function. (A) Multiple sequence alignments and accompanying sequence logo mapped to TgAC10^658-700^. Conserved residues are highlighted by class (blue: hydrophobic, purple: acidic, red: basic, green: polar, orange: Gly, yellow: Pro). (B) Y2H showing interaction of full-length AC9 with the AC10^AC9-BD^ (residues 651-683). (C) Diagram of AC10^Δ(AC9-BD)^ with residues 651-683 deleted from the AC10^wt^ construct. (D) AC10^Δ(AC9-BD)^ localizes properly to the apical cap −/+ IAA. Green, rabbit anti-HA; magenta, mouse anti-V5. (E) Plaque assays show that AC10^Δ(AC9-BD)^-complemented parasites cannot form plaques upon knockdown of endogenous AC10^AID-3xHA^. (F) AC10^Δ(AC9-BD)^ cannot rescue the recruitment of either AC9 or ERK7 to the apical cap. Green, rabbit anti-Myc; magenta, mouse anti-Ty. (G) IFA shows that SAS6L cannot be restored when complemented with AC10^Δ(AC9-BD)^. Green, rabbit anti-V5; magenta, mouse anti-SAS6L. All scale bars are 2 µm.

### A third AC9 binding site on AC10 is required for full parasite fitness

While AC10^AC9-BD^ was sufficient to bind AC9 in our Y2H assay (Fig. 6B), AC10^B^ also contains the second predicted CC domain spanning residues 781-830 (Fig. 1B, 7A). To assess the importance of CC2, we first generated a construct with the AC9-BD deleted from AC10^B^ (AC10^684-1300^) and found that this region still interacted with AC9 (Fig. 7A). We then deleted CC2 from AC10^684-1300^ (AC10^684-1300,ΔCC2^), which resulted in a somewhat attenuated interaction with AC9 in our Y2H assay. We additionally found that a portion of AC10^B^ containing CC2 (AC10^684-913^) is not sufficient for interacting with AC9. These Y2H results suggest that CC2 may contain minor AC9 binding regions and that the remaining residues in AC10^B^ likely provide additional binding sites, further supporting the hypothesis that AC9 and AC10 interact via multiple contact points.

**Fig 7.**
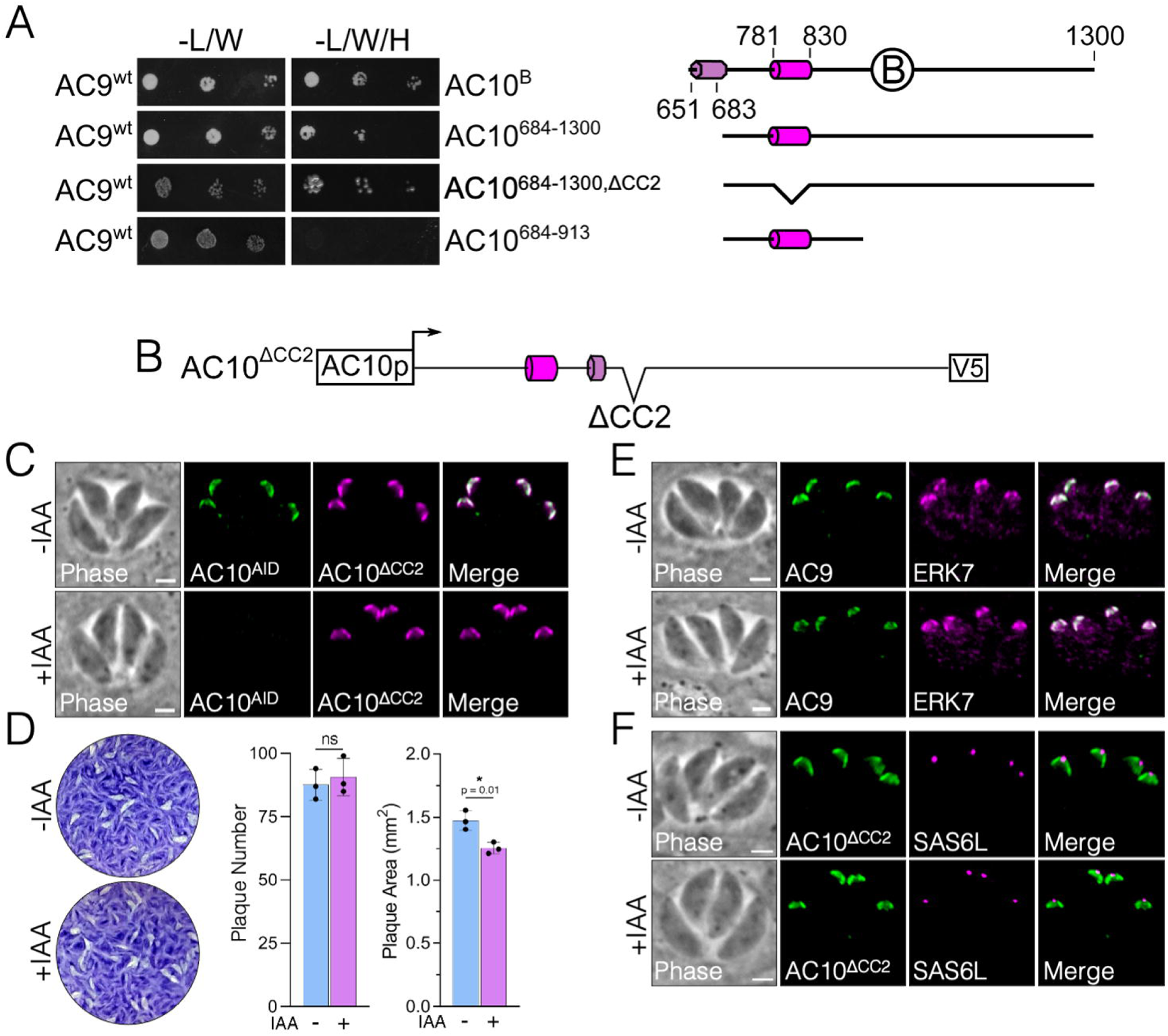
Deletion of CC2 within AC10 results in subtle plaque defects. (A) Y2H to assess interaction of full-length AC9 with the indicated AC10 mutants. Corresponding diagrams of AC10 deletion constructs are shown. (B) Diagram of AC10^ΔCC2^ with residues 781-830 deleted from the AC10^wt^ construct. (C) IFA shows that AC10^ΔCC2^ localizes to the apical cap and is not affected by AC10^AID-3xHA^ knockdown. Green, rabbit anti-HA; magenta, mouse anti-V5. (D) Plaque assays indicate that AC10^ΔCC2^ complementation does not fully rescue the growth defect (15% reduction). Statistical significance was calculated using two-sample two-tailed t tests and p-values are noted on the graph. (E) AC9 and ERK7 staining −/+ IAA shows that AC10^ΔCC2^ can still recruit members of the complex to the apical cap. Green, rabbit anti-Myc; magenta, mouse anti-Ty. (F) IFA illustrates that AC10^ΔCC2^ restores SAS6L staining at the conoid. Green, rabbit anti-V5; magenta, mouse anti-SAS6L. All scale bars are 2 µm.

We then asked whether deletion of CC2 in the context of an otherwise full-length protein would affect AC10 function in parasites. We generated AC10^ΔCC2^ (residues Δ781-830) and expressed it in the triple-tagged AC10^AID-3xHA^ line (Fig. 7B). As with our other deletion constructs, AC10^ΔCC2^ protein localized correctly to the apical cap (Fig. 7C). Upon degradation of AC10^AID-3xHA^, AC10^ΔCC2^ mostly rescued parasite fitness in a plaque assay, with a small but reproducible 15% reduction in plaque size (Fig. 7D). Consistent with this minor impact on the lytic cycle, both AC9 and ERK7 localizations were unaffected (Fig. 7E) and the conoid appeared intact (Fig. 7F). These data suggest that binding of AC9 and other potential interactors at this site, while not required for full parasite fitness, is still functionally relevant.

### AC10 N- and C-terminal deletions reveal additional domains for full apical cap function

The functional regions of AC10 described above only occupy about half of the 1979-residue protein. Notably, AC10 orthologs in other Sarcocystidae are of varying length and display low sequence identity through the majority of the protein (Fig. 1B). To determine if the remainder of the protein harbored any additional regions important for function, we first deleted the N-terminal region of AC10 up to 36 residues N-terminal to AC10^CC1^ (residues 387-1979, AC10^ΔN-term^, Fig. 8A). The AC10^ΔN-term^ protein localized properly to the apical cap independently of AC10^AID-3xHA^ degradation (Fig. 8B). Upon AC10^AID-3xHA^ depletion, parasites with AC10^ΔN-term^ displayed a substantial fitness defect by plaque assay (48% reduction in plaque size, Fig. 8C). However, AC10^ΔN-term^ appears to be sufficient for recruiting both AC9 and ERK7 to the apical cap (Fig. 8D), resulting in the presence of a conoid as demonstrated by apical SAS6L staining (Fig. 8E). Thus, while this N-terminal region is not strictly required for recruiting AC9:ERK7 and maturation of the conoid, its deletion reduces parasite fitness, indicating that this region is important for full AC10 function.

**Fig 8.**
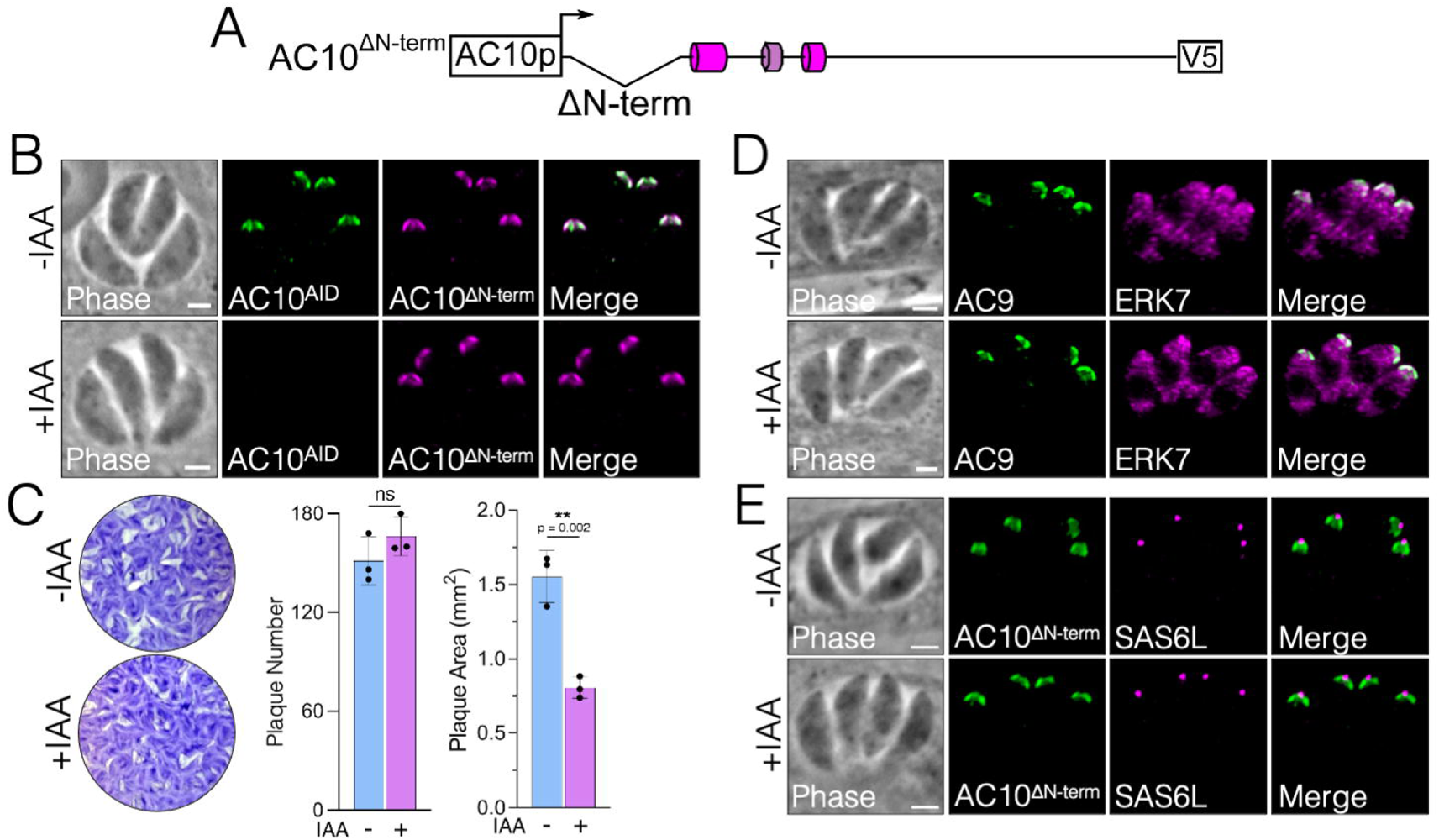
N-terminal deletion of AC10 results in a substantial plaque defect. (A) Diagram of AC10^ΔN-term^ with residues 2-386 deleted from the AC10^wt^ construct. (B) IFA shows that AC10^ΔN-term^ targets properly to the apical cap −/+ IAA. Green, rabbit anti-HA; magenta, mouse anti-V5. (C) Plaque assays show that AC10^ΔN-term^ partially rescues the growth defect, resulting in smaller plaques upon AC10^AID-3xHA^ knockdown (48% reduction). Statistical significance was calculated using two-sample two-tailed t tests and p-values are noted on the graph. (D) IFA illustrates that AC9 and ERK7 are present in the apical cap −/+ IAA. Green, rabbit anti-Myc; magenta, mouse anti-Ty. (E) SAS6L staining indicates that the conoid is present −/+ IAA. Green, rabbit anti-V5; magenta, mouse anti-SAS6L. All scale bars are 2 µm.

We next focused on the C-terminal region of AC10. Due to the lack of identifiable features in this region, we deleted the C-terminal half of the protein, which includes AC10^C^ plus the portion of AC10^B^ C-terminal to the CC domains (residues Δ914-1979, AC10^ΔC-term^, Fig. 9A). Upon examining the localization of AC10^ΔC-term^, we noticed striking, cell-cycle dependent variation. In mature parasites, AC10^ΔC-term^ localized to the apical cap regardless of AC10^AID-3xHA^ depletion (Fig. 9B). However, in budding parasites, AC10^ΔC-term^ was largely absent in the maternal apical cap while remaining intact in the daughter buds (Fig. 9C). We thus assessed the localization of AC9 and ERK7 in mature parasites expressing AC10^ΔC-term^ and found that only a small amount of AC9 could be detected in the apical cap upon AC10^AID-3xHA^ knockdown (Fig. 9D). ERK7 also appeared to be dramatically diminished from the apical cap in mature parasites (Fig. 9D). In budding parasites, while both AC9 and ERK7 were drastically reduced in mature apical caps, the signal appeared largely intact in daughter buds, similar to the localization of AC10^ΔC-term^ (Fig. 9E). Somewhat surprisingly, despite these substantial localization defects, the conoid still appeared to be intact by SAS6L staining, suggesting that the amounts of AC9, AC10^ΔC-term^, and ERK7 in the apical cap are sufficient to stabilize the conoid (Fig. 9F). Nevertheless, plaque assays revealed that parasites expressing AC10^ΔC-term^ suffered a severe defect in parasite fitness upon AC10^AID-3xHA^ degradation (85% reduction in plaque size; Fig. 9G).

**Fig 9.**
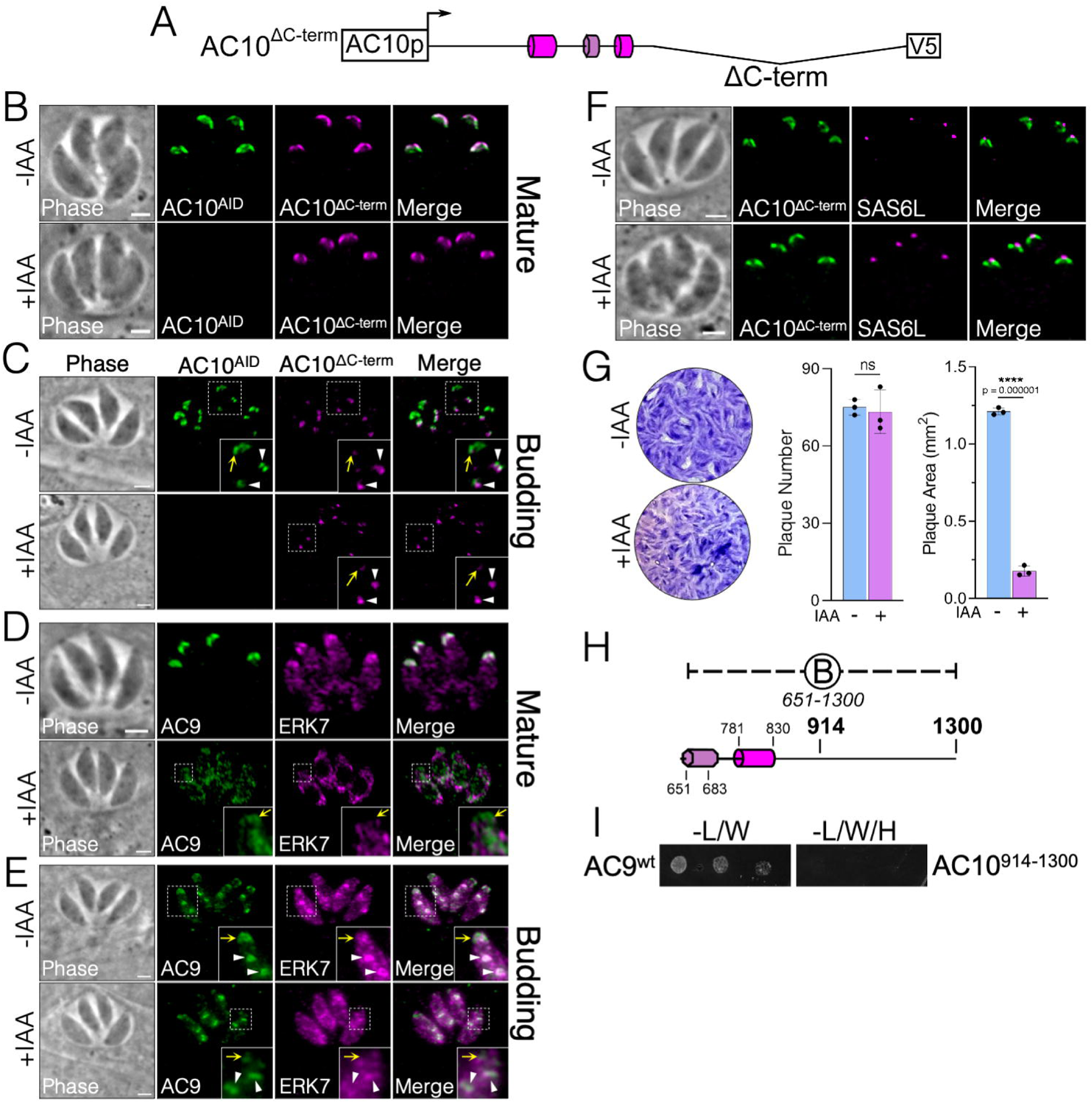
C-terminal deletion of AC10 diminishes maternal apical cap localization and causes severe fitness defects. (A) Diagram of AC10^ΔC-term^ with residues 914-1979 deleted from the AC10^wt^ construct. (B) IFA shows that mature parasites have proper AC10^ΔC-term^ localization −/+ IAA. Green, rabbit anti-HA; magenta, mouse anti-V5. (C) In contrast, actively budding parasites have substantially diminished AC10^ΔC-term^ localization in maternal apical caps (inset, yellow arrows), while AC10^ΔC-term^ localization to daughter buds is unaffected (inset, white arrowheads). Green, rabbit anti-HA; magenta, mouse anti-V5. (D) IFA depicts that both AC9 and ERK7 are substantially mislocalized to the cytoplasm in mature parasites +IAA (insets, yellow arrows). Green, rabbit anti-Myc; magenta, mouse anti-Ty. (E) In budding parasites, IFAs show severely decreased levels of AC9 in the maternal apical cap (insets, yellow arrows) but intact localization in daughter buds (insets, white arrowheads). ERK7 appears absent from the apical cap upon depletion of AC10^AID-3xHA^. Green, rabbit anti-Myc; magenta, mouse anti-Ty. (F) IFAs demonstrate that SAS6L staining appears intact upon degradation of AC10^AID-3xHA^. Green, rabbit anti-V5; magenta, mouse anti-SAS6L. All scale bars are 2 µm. (G) Plaque assays show extremely small plaques upon knockdown of AC10^AID-3xHA^ (85% reduction). Statistical significance was calculated using two-sample two-tailed t tests and p-values are noted on the graph. (H) Diagram illustrating the AC10^914-1300^ construct used in the following Y2H assay. (I) Y2H to assess the interaction of full-length AC9 with AC10^914-1300^.

We next sought to determine whether the C-terminal half of AC10 described above binds directly to AC9. We created a Y2H construct spanning AC10 residues 914-1300 to interrogate the C-terminal portion of AC10^B^ (AC10^914-1300^, Fig. 9H). Despite the defects in AC9 and ERK7 recruitment in AC10^ΔC-term^ parasites, we found that neither AC10^914-1300^ (Fig. 9I) nor the remainder of the AC10 C-terminus (AC10^C^) interacts with AC9 (Fig. 3J). Together, these results suggest that while the AC10 C-terminus does not directly interact with AC9, it contains important regions for maintaining the integrity of the AC9:AC10:ERK7 complex.

Since deletion of either the N- or C-termini of AC10 only partially disrupted function, we assessed whether the combination of these regions is essential by deleting both regions simultaneously (residues Δ2-337 and Δ914-1979, AC10^ΔN/C^, Fig. 10A). As with AC10^ΔC-term^, AC10^ΔN/C^ localized properly in mature parasites (Fig. 10B), and during replication, the signal was diminished specifically in maternal apical caps upon addition of auxin (Fig. 10C). Unlike AC10^ΔC-term^, however, this construct could not rescue the plaque defect at all (Fig. 10D). Western blot analysis demonstrated that the difference between AC10^ΔC-term^ and AC10^ΔN/C^ does not appear to be due to expression levels (Fig. S3B). Consistent with the complete loss-of-function of AC10^ΔN/C^, both AC9 and ERK7 were absent from the maternal apical caps of both mature and budding parasites (Fig. 10E and F). In addition, we observed reduced AC9 and ERK7 signal in the apical caps of daughter buds (Fig. 10F). In agreement with the lack of ability to form plaques, AC10^ΔN/C^ parasites were completely missing apical SAS6L staining upon AC10^AID-3xHA^ depletion (Fig. 10G). Together, these results demonstrate that the cumulative effect of deleting both N- and C-terminal regions renders AC10 nonfunctional.

**Fig 10.**
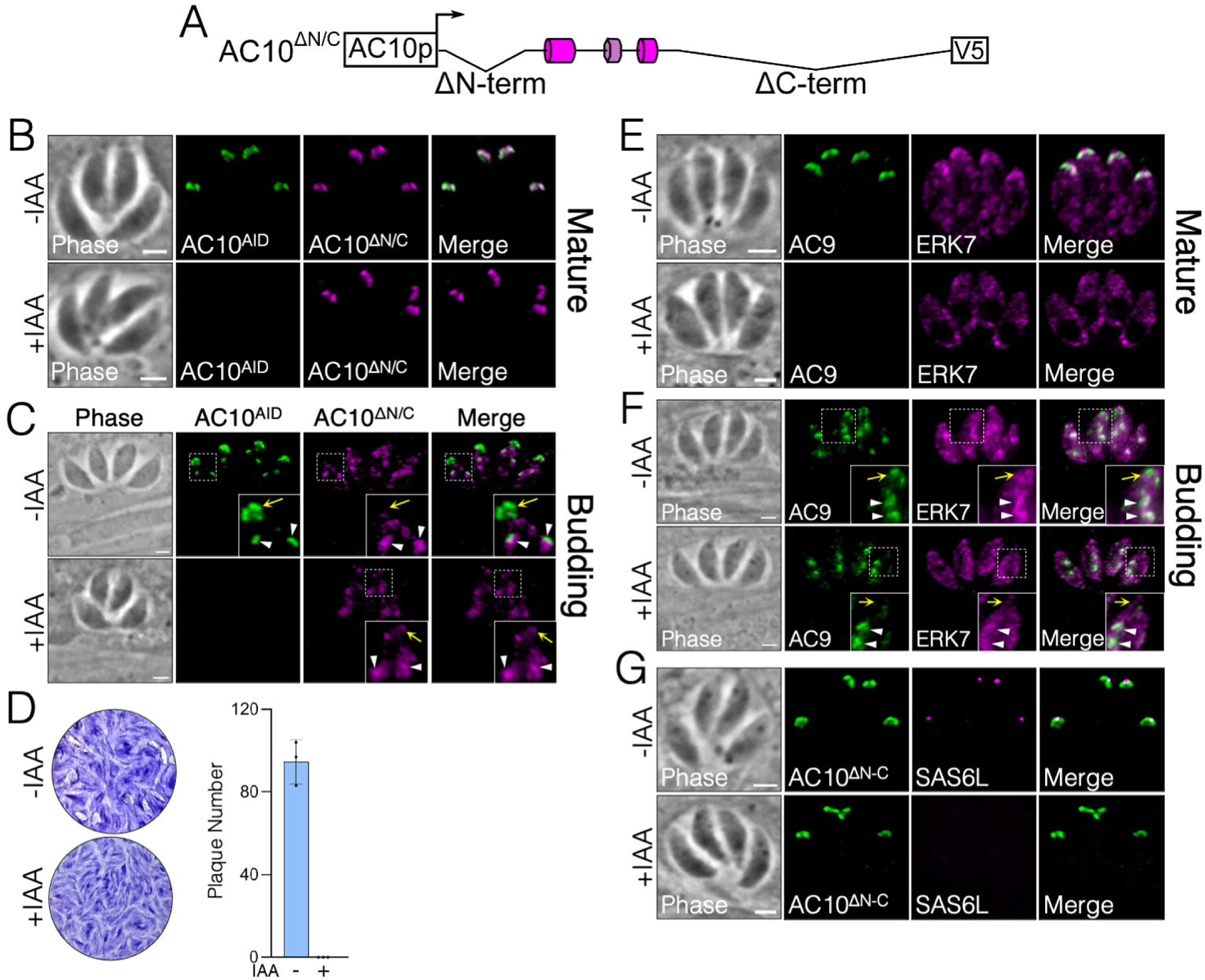
Combination of N- and C-terminal deletions is essential for apical cap function. (A) Diagram of AC10^ΔN/C^ combining the AC10^ΔN-term^ (residues 2-337) and AC10^ΔC-term^ (residues 914-1979) deletions from the AC10^wt^ construct. (B) IFAs illustrate that AC10^ΔN/C^ localizes properly to the apical caps in mature parasites −/+ IAA. Green, rabbit anti-HA; magenta, mouse anti-V5. (C) IFAs show that AC10^ΔN/C^ appears almost completely absent in the maternal apical cap of budding parasites upon depletion of AC10^AID-3xHA^ (insets, yellow arrows). However, in daughter apical caps, AC10^ΔN/C^ remains intact even upon depletion of endogenous AC10^AID-3xHA^ (insets, white arrowheads). Green, rabbit anti-HA; magenta, mouse anti-V5. (D) Plaque assays show that deleting both N- and C-terminal regions from AC10 eliminates plaque formation. (E) IFAs display the absence of AC9 and ERK7 from mature apical caps. Green, rabbit anti-Myc; magenta, mouse anti-Ty. (F) IFAs show that AC9 and ERK7 remain intact in daughter apical caps (insets, white arrowheads) but appear completely eliminated from maternal apical caps upon knockdown of AC10^AID-3xHA^ (insets, yellow arrows). Green, rabbit anti-Myc; magenta, mouse anti-Ty. (G) IFAs display absence of SAS6L upon AC10^AID-3xHA^ knockdown. Green, rabbit anti-V5; magenta, mouse anti-SAS6L. All scale bars are 2 µm.

### AC10 effectively competes with AC9 as an ERK7 substrate

Because AC10 binds both AC9 and ERK7 (Fig. 4, Fig. 5, Fig 6), and ERK7 localization (23) and kinase activity (21) are both essential for a functional conoid, we asked whether AC10 may be phosphorylated by ERK7. Notably, AC10 has 396 phosphorylatable residues (Ser/Thr) and 57 of these residues have been identified as phosphorylated in parasites in published phosphoproteomics datasets (47), including 10 high probability MAPK sites spread throughout the AC10 sequence. We created a bacterial expression construct of the N-terminal region of AC10 that is bound by both AC9 and ERK7. We found that this recombinantly expressed and purified AC10 was robustly phosphorylated by ERK7 (Fig. 11). Remarkably, the AC10 protein was phosphorylated to a much greater degree than myelin basic protein (MBP), a typical generic substrate used to test MAPK activity (48).

**Fig 11.**
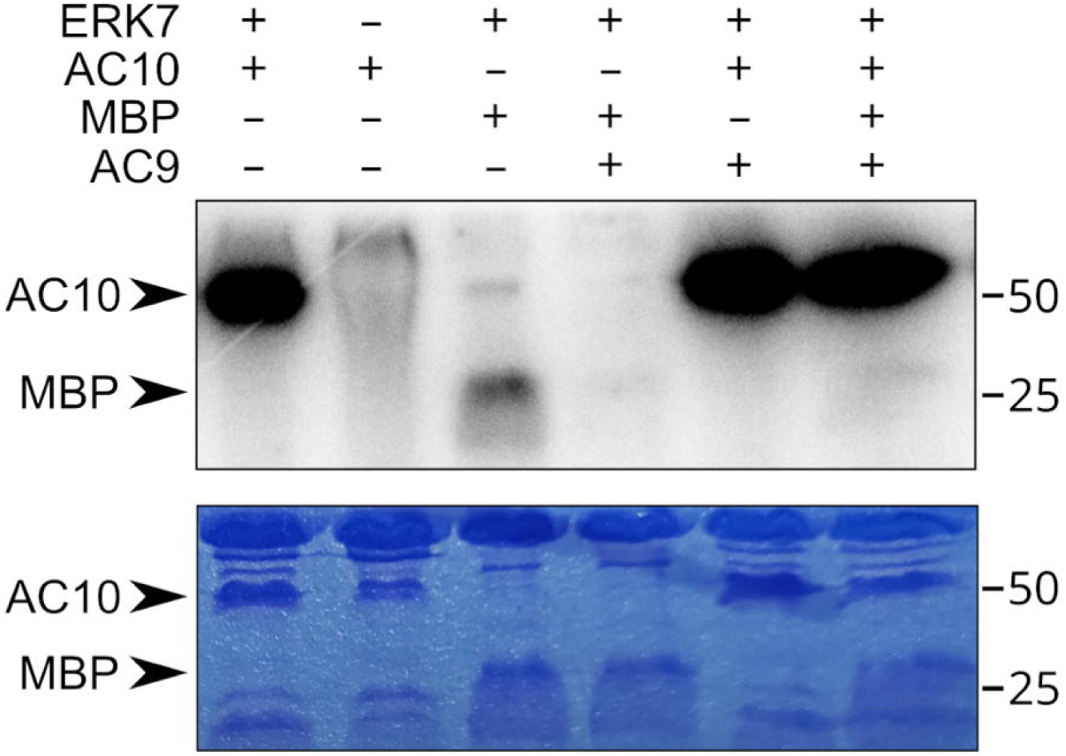
ERK7 robustly phosphorylates AC10 *in vitro.* Autoradiogram and corresponding coomassie stained gel of an *in vitro* kinase assay in which 1 μM ERK7 was used to phosphorylate 10 μM AC10^313-569^ or the generic substrate MBP. In the rightmost 3 lanes, 10 μM inhibitory AC9^418-452^ was added to the reaction. Note that the rightmost lane contains both MBP and AC10 as substrates.

We previously demonstrated that AC9 binds ERK7 with an approximate 20 nM K_D_ and robustly inhibits ERK7 activity (23). This led us to propose a model by which AC9 increases the specificity of ERK7 for its substrates, as true substrates must not only bind the active site, but also compete with AC9 for scaffolding interaction. We therefore tested whether the AC10 interaction with ERK7 is able to overcome inhibition by the AC9^418-452^ peptide (Fig. 11). As expected, addition of equimolar AC9^418-452^ to the kinase reaction completely blocks MBP phosphorylation by ERK7. We found, however, that AC10 phosphorylation is undiminished by the addition of AC9. Furthermore, when we included equimolar AC9, AC10, and MBP in the kinase reaction, we saw that MBP phosphorylation was still fully inhibited, while AC10 was still robustly phosphorylated. These data strongly suggest AC10 is a legitimate substrate of ERK7, and that one function of ERK7 kinase activity may be to regulate the conformation and assembly of the AC10 complex.

## DISCUSSION

In this study, we explore the organization and function of the AC9:AC10:ERK7 ternary complex. We demonstrated that both AC9 and ERK7 are dependent on AC10 to be recruited to the apical cap, suggesting that AC10 is an anchor for the complex. However, it remains unclear how AC10 itself is targeted to the apical cap. One possibility is that other apical cap proteins recruit AC10. Similar to AC10, six of the known apical cap proteins (AC2, AC3, AC4, AC5, AC7, AC8) are associated with the IMC cytoskeletal network (15). Unlike AC9 and AC10, these other apical cap proteins were predicted to be dispensable based on a genome-wide CRISPR screen (49). Thus, it is possible that these apical cap proteins play redundant roles in organizing the AC9:AC10:ERK7 complex. It is also possible that there are undiscovered components of this protein complex or ones that serve to tether AC10 to the apical cap.

To determine how AC9, AC10, and ERK7 interact, we focused on identifiable domains using a combination of pairwise Y2H (Table 1) and complementation assays to assess direct binding and functional relevance. AC10 appears to recruit AC9 (Fig. 2) (22), which in turn recruits ERK7 through a conserved C-terminal motif that serves to both concentrate ERK7 at the apical cap and regulate its kinase activity (23). Our Y2H and complementation assays revealed a conserved helical sequence at the AC9 N-terminus that was both necessary and sufficient to bind AC10 and was required for AC9’s localization at the apical cap (Fig. 2). Remarkably, this single region of AC9 was able to bind multiple sites on AC10 (Fig. 3–7). In addition, AC10 can independently interact with both the kinase domain and C-terminal regions of ERK7 (Fig. 5). AC10 therefore seems to be act as a large scaffolding molecule that recruits multiple copies of each AC9 and ERK7. Furthermore, combined with the multiple binding sites on AC10 for both AC9 and ERK7, because each component of the AC9:AC10:ERK7 complex can interact with the other, it seems likely that AC10 functions to nucleate oligomerization of this complex (Fig. 12). Importantly AC9, AC10, and ERK7 have each been demonstrated to fractionate with the detergent-insoluble parasite cytoskeleton (21, 22), and their oligomerization is consistent with the characteristic meshwork of the IMC cytoskeleton. The AC10 binding region of AC9 is a predicted coiled-coil (AC9^CC^), and we identified two regions of AC10 (AC10^CC1^ and AC10^AC9-BD^) with coiled-coil-like properties that are required for AC9 interaction and essential for AC10 function in parasites. Notably, other predicted coiled-coil domains are essential in other IMC proteins (44, 45, 50), suggesting this may be a general theme of IMC cytoskeleton assembly.

**Fig 12.**
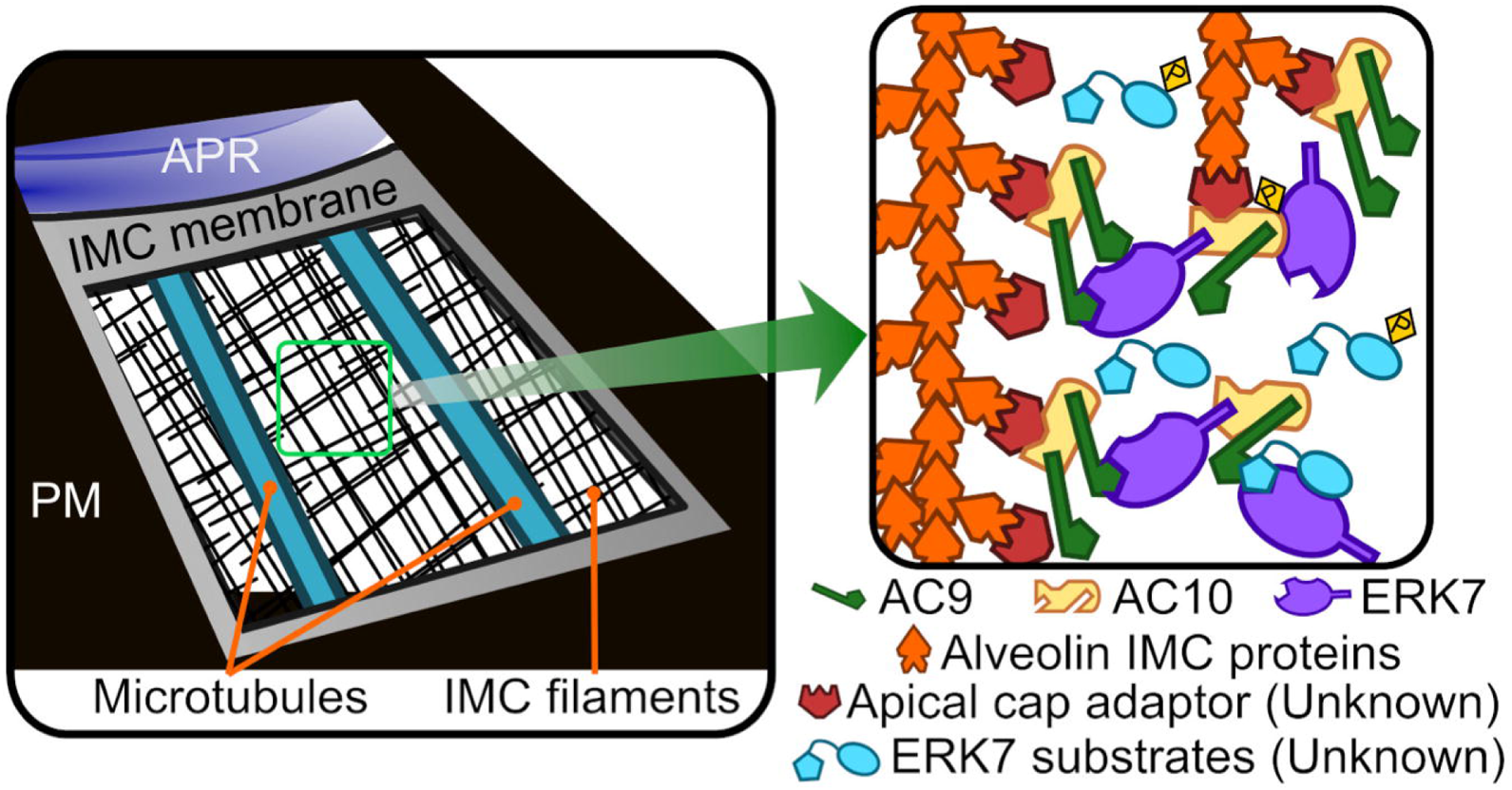
Model for AC9:AC10:ERK7 complex oligomerization in the apical cap. AC9, AC10, and ERK7 oligomerize with the IMC cytoskeleton filaments that are associated with the cytosolic leaflet of the IMC membrane. AC10 recruits the other two proteins to the IMC, possibly through interaction with an undescribed adaptor protein. Because AC10 has multiple binding sites for both AC9 and ERK7, which also interact with one another, the three proteins likely form an irregular oligomer. These interactions concentrate ERK7 at the apical cap while allowing it to bind and phosphorylate its substrates and thereby facilitate the stability of the apical complex.

Deletion of the short AC10^AC9-BD^ sequence blocks AC9 recruitment to the apical cap in parasites (Fig. 6). However, AC9 localization was largely unperturbed in AC10^ΔCC1^ parasites while ERK7 was unable to be recruited to the apical cap (Fig. 4). Remarkably, Y2H revealed that the N-terminal third of AC10 was able to physically interact with both AC9 and the ERK7 kinase domain, though the AC10^CC1^ region itself was only required for AC9 binding (Fig. 4A, Fig. 5). This differential effect of AC10^ΔCC1^ on AC9 and ERK7 binding to this region suggests that the binding interfaces may occupy different surfaces of a folded domain. We also found that this N-terminal region of AC10 was robustly phosphorylated by ERK7 *in vitro* and was unaffected by AC9 inhibition (Fig. 11). Together, these data indicate that AC10 is an ERK7 substrate in parasites and suggest that its phosphorylation functions in regulating the assembly of the AC9:AC10:ERK7 complex into the apical cap cytoskeleton.

While AC10 is found throughout coccidia, its length and much of its sequence are not well conserved (Fig. 1B). Nevertheless, there are stretches of conserved sequence in the N- and C-terminal regions that are outside of those we identified as critical for interacting with AC9 and ERK7. We found that neither of these regions of AC10 were essential to function, though deletion of either reduced parasite fitness (Fig. 8C, 9G). Notably, AC10^ΔC-term^ parasites showed a fragility of the AC9:AC10:ERK7 complex, in which the initial recruitment to the apical cap was largely unaffected in daughter cells (Fig. 9C), but the complex appeared disrupted in mature parasites (Fig. 9D, E). While AC10^ΔC-term^ parasites showed a substantial loss of function, the complex was still able to function in facilitating maturation of the conoid (Fig. 9G). In contrast, deletion of both the N- and C-terminal regions of AC10 rendered the AC9:AC10:ERK7 complex non-functional, as the daughter conoids were lost (Fig. 10G) and parasites were nonviable (Fig. 10D). Therefore, it appears that these regions of AC10 either recruit other, undescribed components of the apical cap cytoskeleton, or form nonessential interactions that facilitate AC9:AC10:ERK7 oligomerization.

This study builds on an increasingly robust body of evidence that the apical cap acts as an essential platform to facilitate the assembly and maintenance of the apical complex (21–23). A previously proposed model suggested that AC9 and AC10 act primarily to stabilize the *Toxoplasma* subpellicular microtubules due to the distribution of AC9 and AC10 proteins along the longitudinal rows of the microtubules (22). While our data support the idea that AC9 and AC10 form filaments in the apical cap cytoskeleton, this model was developed prior to establishing a connection with the MAP kinase ERK7 and its essential role in apical complex maturation (21). We have previously shown that an AC9 mutant that is unable to recruit ERK7 to the apical cap cannot rescue the AC9 knockdown (23). We have built upon that finding here, demonstrating a loss of the conoid in mutant AC10 parasites that are able to recruit AC9, but not ERK7, to the apical cap (Fig. 4J, K). Taken together, our data suggests a different model, in which the ERK7-dependent phosphorylation of AC10 promotes functional assembly of the AC9:AC10:ERK7 complex at the apical cap (Fig. 12). It is likely that ERK7 then phosphorylates other substrates after being recruited at this site, which may include critical components of the apical complex.

## MATERIALS AND METHODS

### *T. gondii* and host cell culture

*T. gondii* RHΔ*ku80*Δ*hxgprt* (parental) and subsequent strains were grown on confluent monolayers of human foreskin fibroblasts (HFFs, ATCC) at 37°C and 5% CO_2_ in Dulbecco’s Modified Eagle Medium (DMEM) supplemented with 5% fetal bovine serum (Gibco), 5% Cosmic calf serum (Hyclone), and 1x penicillin-streptomycin-L-glutamine (Gibco). Constructs containing selectable markers were selected using 1 μM pyrimethamine (dihydrofolate reductase-thymidylate synthase [DHFR-TS]), 50 μg/mL mycophenolic acid-xanthine (HXGPRT), or 40 μM chloramphenicol (CAT) (51–53). Removal of HXGPRT was negatively selected using 350 µg/mL 6-thioxanthine (6-TX), and homologous recombination to the UPRT locus was negatively selected using 5 μM 5-fluorodeoxyuridine (FUDR) (43).

### Antibodies

The HA epitope was detected with mouse monoclonal antibody (mAb) HA.11 (diluted 1:1000) (BioLegend, item no. 901515) or rabbit polyclonal antibody (pAb) anti-HA (diluted 1:1000) (Invitrogen, catalog no. PI715500). The Ty1 epitope was detected with mouse mAb BB2 (diluted 1:1000) (54). The c-Myc epitope was detected with mouse mAb 9E10 (diluted 1:1000) (55) or rabbit pAb anti-Myc (diluted 1:1000) (Invitrogen, catalog no. PA1981). The V5 epitope was detected with mouse mAb anti-V5 (diluted 1:1000) (Invitrogen, catalong no. R96025). *Toxoplasma*-specific antibodies include mouse mAb m-IMC1 (diluted 1:500) (56), mouse mAb anti-ISP1 (diluted 1:1000) (57), rabbit pAb anti-IMC6 (diluted 1:2000) (44).

### Production of IMC12 antibody

The IMC12 coding sequence was cloned into the pET His6 TEV LIC bacterial expression vector (Scott Gradia, Addgene plasmid #29653) using primers P32-35. The construct was transformed into BL21(DE3) *E*. *coli*, and protein was induced with 1 mM IPTG and purified using Ni-NTA agarose under denaturing conditions as described (58). The sample was then dialyzed into PBS to remove the urea, and rabbit antisera was produced by Cocalico Biologicals.

### Immunofluorescence assay and western blot

Confluent HFF cells were grown on glass coverslips and infected with *T*. *gondii*. After 18–24 hours, the coverslips were fixed with 3.7% formaldehyde in PBS and processed for immunofluorescence (IFA) as described (58). Primary antibodies were detected by species-specific secondary antibodies conjugated to Alexa Fluor 488/594 (ThermoFisher). Coverslips were mounted in Vectashield (Vector Labs, Burlingame, CA), viewed with an Axio Imager.Z1 fluorescent microscope (Zeiss), and processed with ZEN 2.3 software (Zeiss). Processing with the ZEN software included deconvolution as well as adaptation of the magenta pseudocolor from the 594 fluorophore.

For western blot, parasites were lysed in 1x Laemmli sample buffer with 100 mM DTT and boiled at 100°C for 10 minutes. Lysates were resolved by SDS-PAGE and transferred to nitrocellulose membranes, and proteins were detected with the appropriate primary antibody and corresponding secondary antibody conjugated to horseradish peroxidase. Chemiluminescence was induced using the SuperSignal West Pico substrate (Pierce) and imaged on a ChemiDoc XRS+ (Bio-Rad).

### Endogenous epitope tagging

For C-terminal endogenous tagging, a pU6-Universal plasmid containing a protospacer against the 3′ untranslated region (UTR) approximately 100-200 bp downstream of the stop codon was generated for AC9, AC10, and ERK7, as described previously (59). A homology-directed repair (HDR) template was PCR amplified using the LIC vectors p3xHA-mAID.LIC-HXGPRT, p3xMyc.LIC-DHFR, p2xStrep3xTy.LIC-HXGPR that include the epitope tag, 3′ UTR, and a selection cassette (60). The HDR templates include 40 bp of homology immediately upstream of the stop codon or 40 bp of homology within the 3′ UTR downstream of the CRISPR/Cas9 cut site. This template was amplified in 400 μL, purified by phenol-chloroform extraction, ethanol precipitated, and electroporated into RH*ΔhxgprtΔku80* parasites, along with 50 μg of the pU6-Universal plasmid. Successful tagging was confirmed by IFA, and clonal lines of tagged parasites were obtained through limiting dilution. AC10, AC9, and ERK7 were tagged using CRISPR/Cas9 with primers P1-P12. This process was followed to generate the triple-tagged parasites (AC10^AID-3xHA^ | AC9^3xMyc^ | ERK7^3xTy^).

### Complementation of AC9 and AC10

The AC9 wild-type complementation construct (23) was used as the template for creating a deletion of the CC domain. The online NEBasechanger (https://nebasechanger.neb.com/) was used to design primers and the Q5 Site Directed Mutagenesis Kit (NEB) was used to generate pUPRTKO-ISC6pro-AC9^ΔCC^-3xTy (primers P13-14). Both the AC9^wt^ and AC9^ΔCC^ constructs were linearized with DraIII-HF (NEB), transfected into AC9^AID-3xHA^ parasites along with a universal pU6 that targets the UPRT coding region, and selected with 5 μg/mL FUDR for replacement of UPRT as described (43).

For AC10, the endogenous promoter as well as the full coding region was PCR amplified from genomic DNA. This was cloned into the pUPRTKO vector (23) with Gibson assembly (primers P15-18), resulting in pUPRTKO-AC10pro-AC10^wt^-1xV5. The online NEBuilder tool was used to design these Gibson primers (https://nebuilder.neb.com/#!/).

This complementation vector was then linearized with PsiI-v2 (NEB), transfected into triple-tagged parasites, and selected with FUDR. Clones expressing the pUPRTKO-AC10pro-AC10^wt^-1xV5 vector were screened by IFA, and a V5-positive clone was designated AC10^wt^. For most of the AC10 deletion constructs, pUPRTKO-AC10pro-AC10^wt^-1xV5 was used as the template for Q5 Site Directed Mutagenesis Kit (NEB) (primers P19-28). For AC10^ΔN/C^ construct, Gibson assembly was used with pUPRTKO-AC10pro-AC10^wt^-1xV5 as the template for the vector (primers P29-30) and wildtype cDNA was used as a template for the insert (primers P31-32). The same processes for linearization, transfection, and selection as described above were followed for all deletion constructs.

### Plaque assays

Six-well plates with HFF monolayers were infected with equal numbers of individual strains grown −/+ 500 µM IAA. Plaques were allowed to form for 7 days, fixed with ice-cold methanol, and stained with crystal violet. The areas of 30 plaques per condition were measured using ZEN software (Zeiss). All plaque assays were performed in triplicate for each condition. Graphical and statistical analyses were performed using Prism GraphPad 8.0. Multiple two-tailed t-tests were used to compare the SD-centered means of −/+ IAA and statistical significance was determined using the Holm-Sidak method.

### Pairwise yeast-2-hybrid

ERK7 and AC9 sequences were cloned into the pB27 vector (Hybrigenics SA) as N-terminal fusions with the LexA DNA binding domain by Gibson assembly or enzyme inverse mutagenesis. AC10 sequences were cloned into the pP6 vector (Hybrigenics SA) as N-terminal fusions with the GAL4 activating domain. AC9 and AC10 constructs were created by Gibson assembly using *Toxoplasma* expression constructs as template and additional truncations were made by enzyme inverse mutagenesis with primers P36-56. ERK7 truncations were created from a full-length pB27 construct provided by Hybrigenics using primers P57-58. Synthetic dropout media was purchased from Sunrise Science. To test for interactions, pairs of constructs were transformed into the L40 strain of *S. cerevisiae* (MATa his3Δ200trp1-901 leu2-3112 ade2 LYS2::(4lexAop-HIS3) URA3::(8lexAop-lacZ) GAL4; gift of Melanie Cobb). Strains were grown overnight in permissive (-Leu/-Trp) media, normalized to their OD_600_, and spotted in 5x dilution in both permissive and restrictive (-Leu/-Trp/-His) media. Relative growth in the two conditions was assessed after 3-4 days incubation at 30°C.

### Protein expression and purification

All recombinant proteins were expressed as N-terminal fusions to His_6_-SUMO in Rosetta2 (DE3) bacteria overnight at 16°C overnight after induction with 300 mM IPTG. Cells were resuspended in binding buffer (50 mM Tris, pH 8.6, 500 mM NaCl, 15 mM Imidazole) and lysed by sonication. His_6_-tagged protein was affinity purified using NiNTA resin (Qiagen), which was washed with binding buffer. Protein was eluted in 20 mM Tris, pH 8.6, 100 mM NaCl, 150 mM Imidazole. Protein was diluted 1:1 with 20 mM Tris, pH 8.6 and purified by anion exchange on a HiTrapQ column. For ERK7 kinase and AC9^418-452^, anion exchange peaks were pooled, incubated with ULP1 protease for 30 min, after which they were diluted 1:1 in water and the cleaved SUMO separated from the protein of interest by anion exchange. The flow-through was concentrated and purified by size-exclusion chromatography, after which it was flash frozen in 10 mM HEPES, pH 7.0, 300 mM NaCl for storage.

### *In vitro* kinase assay

ERK7 kinase activity was assessed using 1 µM purified ERK7 kinase, 5 mM MgCl_2_, 200 µM cold ATP, 10 mM DTT, 1 mg/mL BSA, 300 mM NaCl, 20 mM HEPES pH 7.0, 10% glycerol. Reactions were started by adding a hot ATP mix that contained 10 µCi ɣ[ 32 P] ATP and 5 µg MBP and/or 10 μM AC10^313-569^ as substrate and in the presence or absence of 10 μM AC9^418-452^. The 25 µL reactions were incubated in a 30°C water bath for 30 min. Reactions were stopped by adding 5 µL 6x SDS-buffer. 10 µL of each reaction was then separated by SDS-PAGE. Gels were fixed and coomassie stained and the extent of phosphorylation was assessed by phosphorimager (GE Typhoon).

## Acknowledgements

We thank Gary Ward for anti-IMC1 antibodies and members of the Reese and Bradley labs for helpful reading of the manuscript.

## FIGURE LEGENDS

**Fig S1. Antibody validation for IMC12.**

(A) IFAs show the IMC12 antibody co-localized with IMC1. Upper panels show mature parasites while the lower panels show ones in the process of budding, highlighting that IMC12 localizes exclusively to the maternal IMC. Green, rabbit anti-IMC12; magenta, mouse anti-IMC1. IFA scale bars are 2 µm. (B) Western blot analysis validates the efficacy of the IMC12 antibody. Endogenously tagged IMC12^3xMyc^ parasites display the upshift in protein size due to the mass of the epitope tag compared to untagged parasites, solidifying the identity of the band detected by the IMC12 antibody. IMC12 detected with rabbit anti-IMC12; IMC12^3xMyc^ detected with mouse anti-Myc. Rabbit anti-IMC6 was used as a loading control.

**Fig S2. Line intensity scans of ERK7 localization.**

Fluorescence intensity was measured across the indicated white lines and the resulting relative intensity values from the four lines were averaged to produce the line intensity graph. Orange shading depicts the approximate position of the apical cap.

**Fig S3. Relative protein expression levels of mislocalized AC9 and AC10.**

(A) Western blot of whole cell lysates showing the protein expression levels of AC9^wt^ and AC9^ΔCC^ −/+ IAA. AC9^wt^ and AC9^ΔCC^ were detected with mouse anti-Ty1 and rabbit anti-IMC12 was used as loading control. (B) Western blot showing migration of the indicated AC10 complementation constructs −/+ IAA. AC10^wt^ undergoes substantial breakdown during processing (also see Fig. 2C). Red arrows indicate the likely primary translation product for each construct. AC10 constructs were detected with mouse anti-V5 and rabbit anti-IMC12 was used as loading control.

**Fig S4. Control Y2H experiments.**

Y2H demonstrating lack of autoactivation of the indicated constructs. Each construct is co-expressed with the corresponding empty bait or prey vectors, as appropriate.

**Table S1. Oligonucleotides used in this study.**

All primer sequences are shown in the 5’ to 3’ orientation.

