## Supplementary figures and images for "Multivalent interactions drive the *Toxoplasma* AC9:AC10:ERK7 complex to concentrate ERK7 in the apical cap"

### Supplemental Figure 1

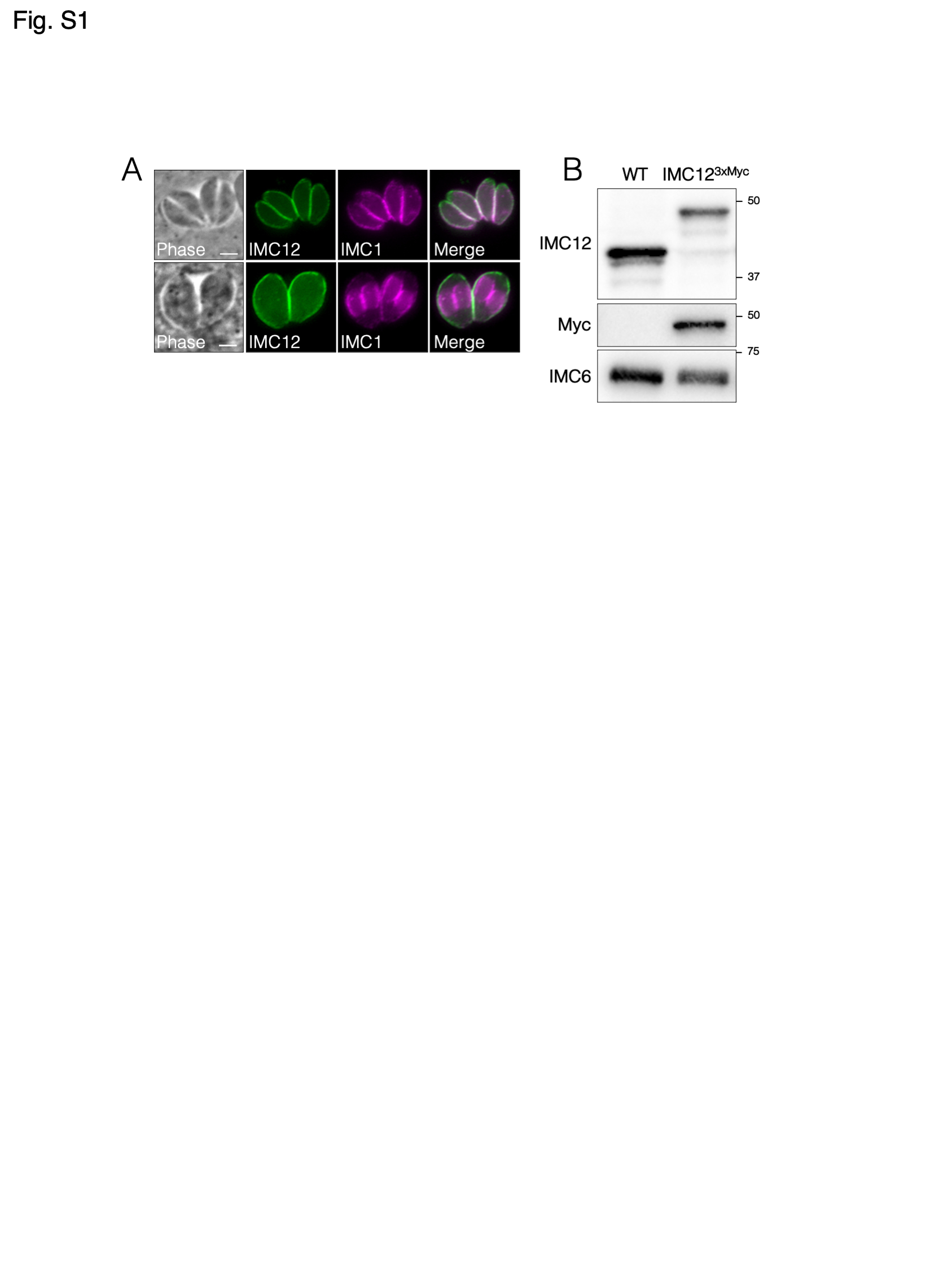

### Supplemental Figure 2

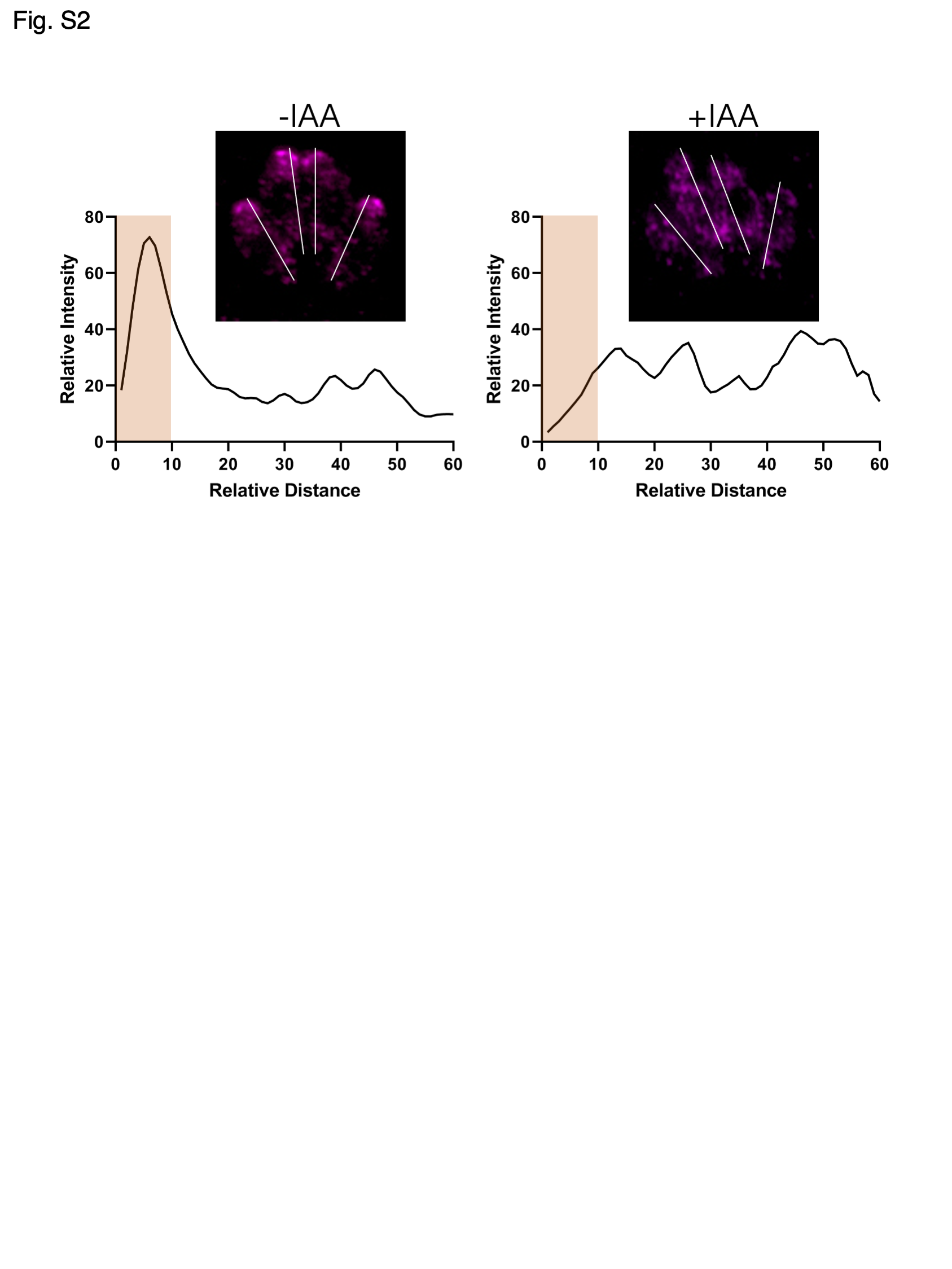

### Supplemental Figure 3

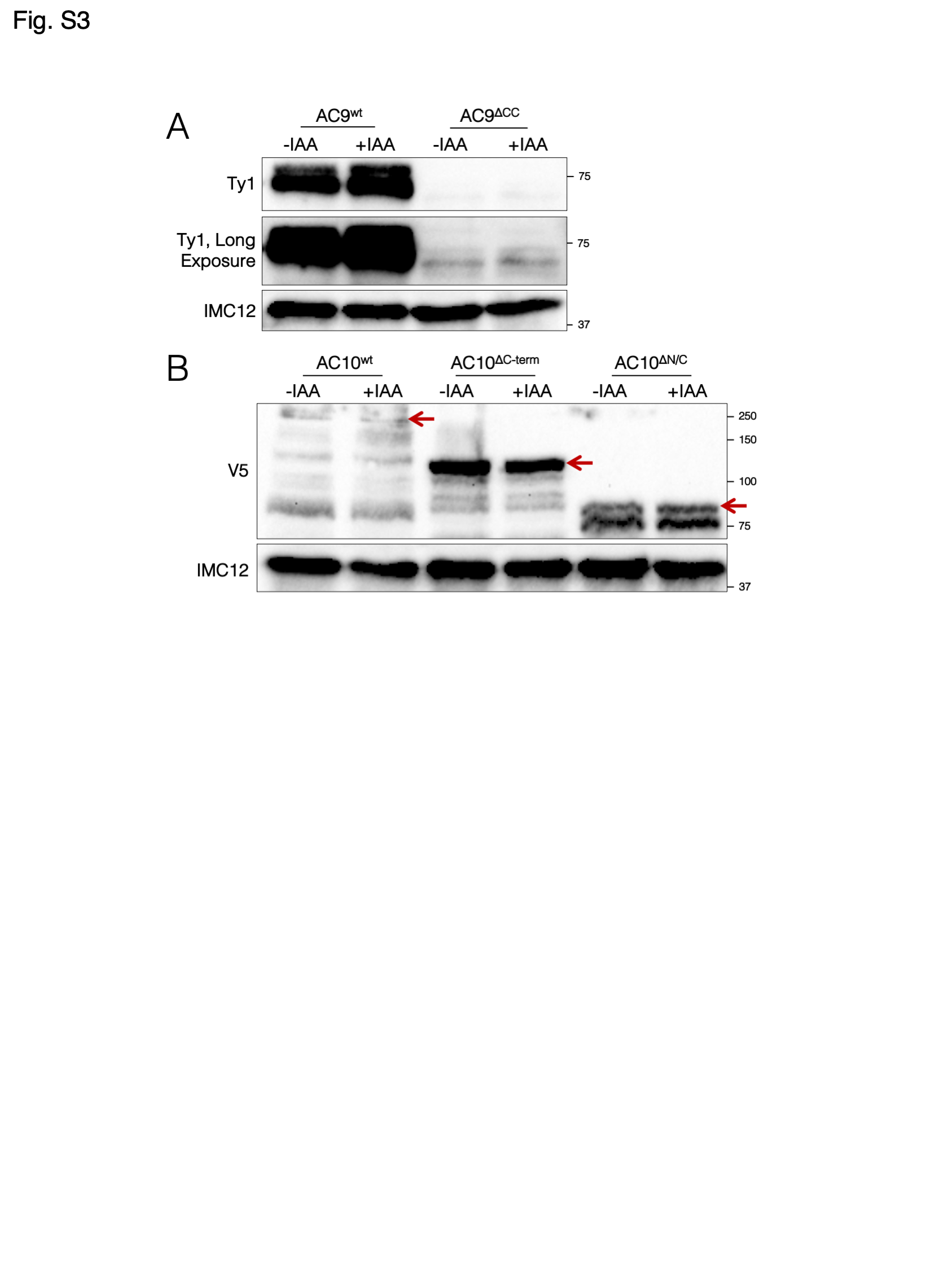

### Supplemental Figure 4

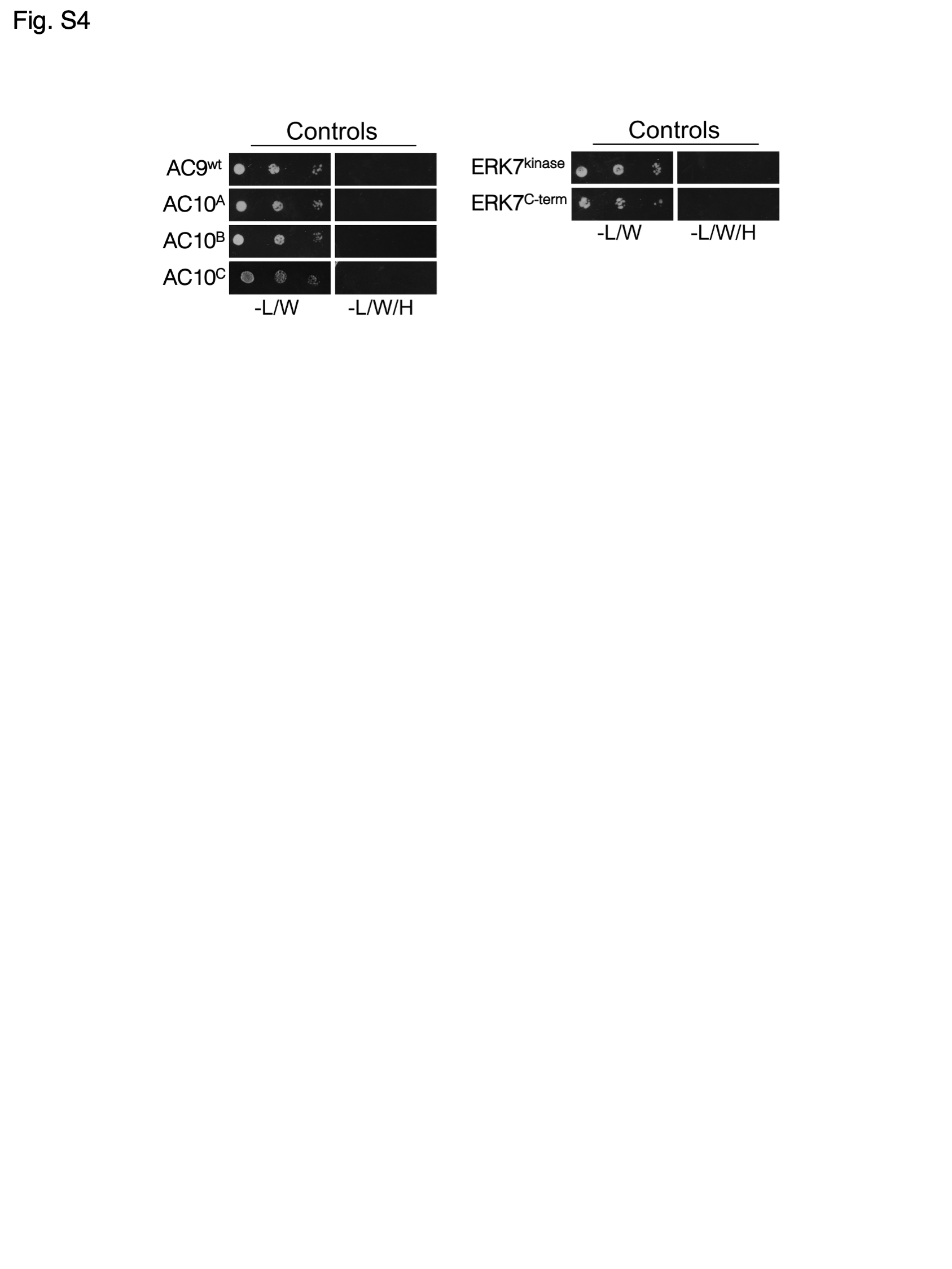
