## Supplemental Table 1 for "Multivalent interactions drive the *Toxoplasma* AC9:AC10:ERK7 complex to concentrate ERK7 in the apical cap"

**Supplemental Table 1: Oligonucleotides used in this study.**

| Purpose | Name | Description | Sequence 5'-3' |
|---|---|---|---|
| **AC10 (TGGT1_292950) tagging** | P1 | AC10 gRNA-tagging sense | AAGTTgTAAGGGTGCAAGAGTTGATGG |
| | P2 | AC10 gRNA-tagging antisense | AAAACCATCAACTCTTGCACCCTTAcA |
| | P3 | AC10 5' HDR template | GCTGAAGACTCGCCTGATAAAGCTCACGAAGGCAAACACGGGAAGTGGAGGACGGGAATT |
| | P4 | AC10 3' HDR template | AGTCCACTGCTGCTCCTCGAGCAGTCTGGGAGATTTGGCGACGGCCAGTGAATTGTAATA |
| **AC9 (TGGT1_246950) tagging** | P5 | AC9 gRNA-tagging sense | AAGTTATGTGTTCCTCAAATGTCAGG |
| | P6 | AC9 gRNA-tagging antisense | AAAACCTGACATTTGAGGAACACATA |
| | P7 | AC9 5' HDR template | GTCGGGGAACCGCGAACCGAGTGAATATCCGCAGGGAATGGGAAGTGGAGGACGGGAATTC |
| | P8 | AC9 3' HDR template | CTCGCAGCGTGTCGCACTGAACTCTTGTGCGGAGAGAGCGACGGCCAGTGAATTGTAATA |
| **ERK7 (TGGT1_233010) tagging** | P9 | ERK7 gRNA-tagging sense | AAGTTGCAAAAGCAAAGATTCAGACG |
| | P10 | ERK7 gRNA-tagging antisense | AAAACGTCTGAATCTTTGCTTTTGCA |
| | P11 | ERK7 5' HDR template | TTCTCTTTTTTTTCAGTCTGCGTCCAAGACATACAACAGCGGAAGTGGAGGACGGGAATT |
| | P12 | ERK7 3' HDR template | GCTTTCTCCACCTTCGCTTTCGGTGGAAGTCTTGCGAGCGACGGCCAGTGAATTGTAATA |
| **AC9 CC deletion construct** | P13 | AC9 CCdel mut fwd | AAGGAGGCGTTTGCACGCAG |
| | P14 | AC9 CCdel mut rev | GGCTGCTGCAGCCGGGTC |
| **AC10 promoter and full length complementation** | P15 | pUPRTKO vector fwd | GCGGCCGCCTACCCGTAC |
| | P16 | pUPRTKO vector rev | ATGCATATGCGATGTCGAACCCTCG |
| | P17 | AC10pro-coding fwd | gttcgacatcgcatatgcatCCCCACTCGTTCCTCGATG |
| | P18 | AC10pro-coding rev | tcgtacgggtaggcggccgcTCTCGCCTTTAATTGCAGTACG |
| **AC10 CC1 deletion construct** | P19 | AC10CC1 Q5del fwd | CTGACGCATGCAGTGGAC |
| | P20 | AC10CC1 Q5del rev | GTCACCGCAGGTGGCATC |
| **AC10 AC9-BD deletion construct** | P21 | AC10-AC9bd Q5del fwd | AAACTAGACGCTGAAGAC |
| | P22 | AC10-AC9bd Q5del rev | GTTATCTTCTGCACTGCC |
| **AC10 CC2 deletion construct** | P23 | AC10CC2 Q5del fwd | GAAAGTACAGGAAGTCGTG |
| | P24 | AC10CC2 Q5del rev | ACAGACATTCGATCGTTG |
| **AC10 N-term deletion construct** | P25 | AC10N-term Q5del fwd | GGAGACACGCGTCCGCAT |
| | P26 | AC10N-term Q5del rev | CATCGCAACTTCCTCCTTTTCG |
| **AC10 C-term deletion construct** | P27 | AC10C-term Q5del fwd | GCGGCCGCCGGCAAACCT |
| | P28 | AC10C-term Q5del rev | GGACTGAGCAGGTAGAACTGAAAGGGATAGAGG |
| **AC10 N/C deletion construct** | P29 | pUPRTKO-AC10wt-vector-fwd | GCGGCCGCCGGCAAACCT |
| | P30 | pUPRTKO-AC10wt-vector-rev | CATCGCAACTTCCTCCTTTTCGCAAAAAGATGTGTTCTTTC |
| | P31 | AC10N-Cdel fwd | aaaaggaggaagttgcgatgCTTACGCGTCTAGATTTG |
| | P32 | AC10N-Cdel rev | ataggtttgccggcggccgcGGACTGAGCAGGTAGAAC |
| **IMC12 pET28 construct for antibody production** | P32 | IMC12-pET28 fwd | TAATAACATTGGAAGTGGATAAC |
| | P33 | IMC12-pET28 rev | TGCATTGGATTGGAAGTAC |
| | P34 | IMC12-coding fwd | tgtacttccaatccaatgcaGCAACCGAGTTCGTCGTTC |
| | P35 | IMC12-coding rev | atccacttccaatgttattaCTGGGGCATGGAGTCGAC |
| **Yeast-2-hybrid** | P36 | f pB27* ga/PhQC [58] | TAAGGGCCACTGGGGCCCC |
| | P37 | r pB27 ga/PhQC [59] | TCCGGCCCCGAATTCCAGC |
| | P38 | f pP6 3p ga/PhQC [57] | CTCGAGTAGCTAGTGTCTAGAG |
| | P39 | r pP6 5p ga/PhQC [57] | ATTCGTGGCCCCTGCGG |
| | P40 | f AC9(D2) pb27-ga [58] | ctggaattcggggccggaGACGTCTCCGGTCGAGGC |
| | P41 | r AC9(M452*) pb27-ga | gggccccagtggcccttaCATTCCCTGCGGATATTCACTCG |
| | P42 | f AC9(P70) blunt [61] | CCGGCTGCAGCAGCCATTC |
| | P43 | r AC9(Q113*) blunt [59] | tcaCTGCTCCTCCAAGCTTTTGAAAG |
| | P44 | r AC9(A157*) blunt [59] | tcaGGCAAAGCCTTGAATGGTTGAGG |
| | P45 | f AC10(V2) pP6-ga [57] | gccgcaggggccacgaat GTGACTGCAGTACCCAATCCTTC |
| | P46 | r AC10(N650*) pP6-ga [57] | ctagacactagctactcgag tca GTTATCTTCTGCACTGCCTTTGAC |
| | P47 | f AC10(E651) pP6-ga [59] | gccgcaggggccacgaatGAAGACAGCACAGAGAGCCAG |
| | P48 | r AC10(T1300*) pP6-ga [59] | ctagacactagctactcgagtcaGGTGGTACGCGTCGCTATTC |
| | P49 | f AC10(S1301) pP6-ga [57] | gccgcaggggccacgaatTCCCCTTTTCAGCGGGGAG |
| | P50 | r AC10(R1979*) pP6-ga [57] | ctagacactagctactcgagTCATCTCGCCTTTAATTGCAGTAC |
| | P51 | f AC10(K684) blunt [59] | AAACTAGACGCTGAAGACCAGAAG |
| | P52 | r AC10(R683) blunt [58] | GCGTGGATCAAGCACCTTGG |
| | P53 | r AC10(S913*) blunt [58/62] | tcaGGACTGAGCAGGTAGAACTG |
| | P54 | f AC10(W914) blunt [59/64] | TGGAATACTACGTCTGTGTCCGC |
| | P55 | r AC10(C780 no*) [57/61] | ACAGACATTCGATCGTTGCTG |
| | P56 | f AC10(E831) [57/61] | GAAAGTACAGGAAGTCGTGCG |
| | P57 | r TgERK7(A358*) blunt [61] | tcaAGCTGTGCGGTGTCGCCG |
| | P58 | f TgERK7(G359) blunt [59] | GGTTCTTCCGGCCGCCACC |
